# Systematic identification of context-dependent gene-essentiality in Glioblastoma: The GBM-CoDE platform

**DOI:** 10.1101/2025.01.27.634260

**Authors:** Mitchell T Foster, Ailith Ewing, Margaret Frame, Paul M Brennan, Ava Khamseh, Sjoerd V Beentjes, Neil O Carragher, Colin A Semple

## Abstract

Glioblastoma (GBM) is a heterogeneous and aggressive brain tumour that is invariably fatal despite maximal treatment. Genetic or transcriptomic ‘biomarkers’ could be used to stratify patients for treatments, however, pairing biomarkers with appropriate therapeutic ‘targets’ is challenging. Consequently, therapeutics have not yet been optimised for specific GBM patient subsets. Here we integrate genome-wide CRISPR/Cas9 knockout screening and genetic-annotation data for 60 distinct patient-derived, IDH^wildtype^, adult GBM cell lines, quantifying the essentiality of 15,145 genes. We describe a novel method using Targeted Learning, to estimate the effect size of GBM-relevant biomarkers on context-dependent gene essentiality (GBM-CoDE). We derive multiple target-biomarker pair hypotheses, which we release in an accessible platform to accelerate translation to biomarker-stratified clinical trials. Two of these (WWTR1 with EGFR mutation/amplification, and VRK1 with VRK2 expression suppression) have been validated in GBM, implying that our additional novel findings may be valid. Our method is readily translatable to other cancers of unmet need.

## Introduction

GBM shows complex heterogeneity within and between patients, but in the absence of effective biomarker-driven therapies, all patients currently receive the same initial treatment after surgery: temozolomide and radiotherapy^1^. Developing therapies that can be positioned towards tumours with specific driver mutations, transcriptomic subtypes, or cell states may overcome the failure of therapies tested in GBM patient populations unselected for such molecular features.

The genetic modifications associated with cellular malignancy can expose therapeutic vulnerabilities. For example, the PARP-inhibitor Olaparib is more effective in cancer cells that have a deficiency in homologous recombination (HRD). HRD can be caused by various genetic abnormalities but is most notably associated with loss-of-function mutations in the tumour suppressor genes BRCA1 and BRCA2^2^. BRCA1 and PARP1 are a ‘synthetically lethal’ gene pair because cells are non-viable when both genes are depleted, but viable if either one is absent in isolation^2,3^. A more inclusive term, ‘context-dependent gene essentiality,’ describes how the fitness defect induced by knockout of a gene can depend on multiple cell-intrinsic biological properties, that are not limited to the function of a single synthetically lethal pair gene. For example, PARP1 is essential in the context of HRD. Here the therapeutic ‘target’ is PARP1, and two possible ‘biomarkers’ for when to therapeutically inhibit its gene product, include either a mutation in BRCA1 or the presence of HRD. We hypothesized that there may be undiscovered context-dependent gene essentiality in GBM (GBM-CoDE), that could enable acceleration towards biomarker-stratified clinical trials. However, there is a pressing need to develop methods to pair therapeutic target genes with the known GBM biomarkers seen clinically.

Here we identify, process, and integrate datasets from multiple sources^4–8^ to arrive at a curated dataset comprising 60 adult IDH-wildtype (IDH^wt^)^9,10^ GBM cell lines that have been subjected to pooled genome-wide CRISPR-Cas9-knockout-mediated screening. We annotated these cell lines with genetic and transcriptomic features that have been described in human GBM^5,11–16,16–22^. We developed a computational method that uses Targeted Learning^23^ to systematically and robustly identify context-dependent gene essentiality, and applied it to these datasets. Targeted Learning - the combination of Super Learning (SL) and Targeted Maximum-Likelihood Estimation (TMLE) - is a mathematical framework for model-independent quantity estimation that benefits from mathematical guarantees of coverage of the ground truth and, consequently, realistic p-values^23^. Using this approach we identify multiple novel gene targets of therapeutic interest for inhibition in GBM in the context of clinically relevant biomarkers. These target-biomarker pair hypotheses, if experimentally validated, could translate to biomarker-stratified clinical trials in GBM. In this manuscript, we describe our method, discuss a subset of our results, and release our source data. Furthermore, we will make all of our results available in a user-friendly software interface to enable other groups to explore novel biomarker strategies in GBM. Finally, our approach is readily repurposable in other cancers of unmet need.

## Results

### An integrated dataset of gene essentiality reveals GBM-specific commonly essential genes

We collected data from multiple studies^4–8^ in which 83 GBM cell lines had been subject to genome-wide pooled single-guide-RNA (sgRNA) library CRISPR/Cas9-mediated knockout screening (Table S1). We performed copy-number correction using CRISPRCleanR^24^ and then used the Bayesian Analysis of Gene Essentiality (BAGEL2) algorithm^25^, with multi-target correction mode enabled, to unite both the effect-size and statistical confidence that a gene is essential, into a single, continuous quantity, the log Bayes Factor (BF) (see Methods). After technical and biological quality-control, and identification of duplicated cell lines between studies, individual cell line datasets were integrated and normalised (see Methods). This process culminated in an integrated dataset comprising 60 distinct, human adult patient-derived IDH^wt^ GBM cell lines in which the essentiality of 15,145 genes were quantified.

Within these GBM cell lines, we identified 650 core-essential genes (genes with a BF ≥ 6 in ≥ 85% of cell lines^26^): a set of genes that are necessary for GBM cell survival, irrespective of the genomic or transcriptomic context (see Methods). As expected, all of these genes were found within a set of ‘generally essential’ genes which we defined: genes that are either essential across multiple cancer types^27^, in human stem cells^28^, or within critical biological processes which are not cell type dependent^29^ (see Methods; Fig 1a). This reiterated the heterogeneity of, and the necessity to identify context-dependent gene essentiality, *within* GBM. We then identified ‘GBM-core-fitness’ genes, and ‘GBM-commonly-essential’ genes using the using Adaptive Daisy Model (ADaM^29^) and the Fitness Percentile (FiPer^29^) algorithms respectively, both of which use a less strict threshold for essentiality (see Methods). Intersecting these, and excluding the ‘generally essential’ genes, we defined a set of 39 ‘GBM-specific commonly essential’ genes that were enriched most notably for involvement in kinase binding, regulation of differentiation and regulation of adhesion-dependent cell spreading (Fig 1; see Methods). Some of the products from genes within this set (such as VRK1, PTK2, WWTR1, GPX4, CDC25B and PKN) are established druggable targets which can be inhibited with existing small molecules. Doing so may theoretically perturb GBM cells, but be less toxic to normal tissue than suppression of ‘generally essential’ gene products. However, because of the heterogeneity seen in GBM, we additionally wanted to identify biomarkers that influenced gene essentiality.

**Fig 1:**
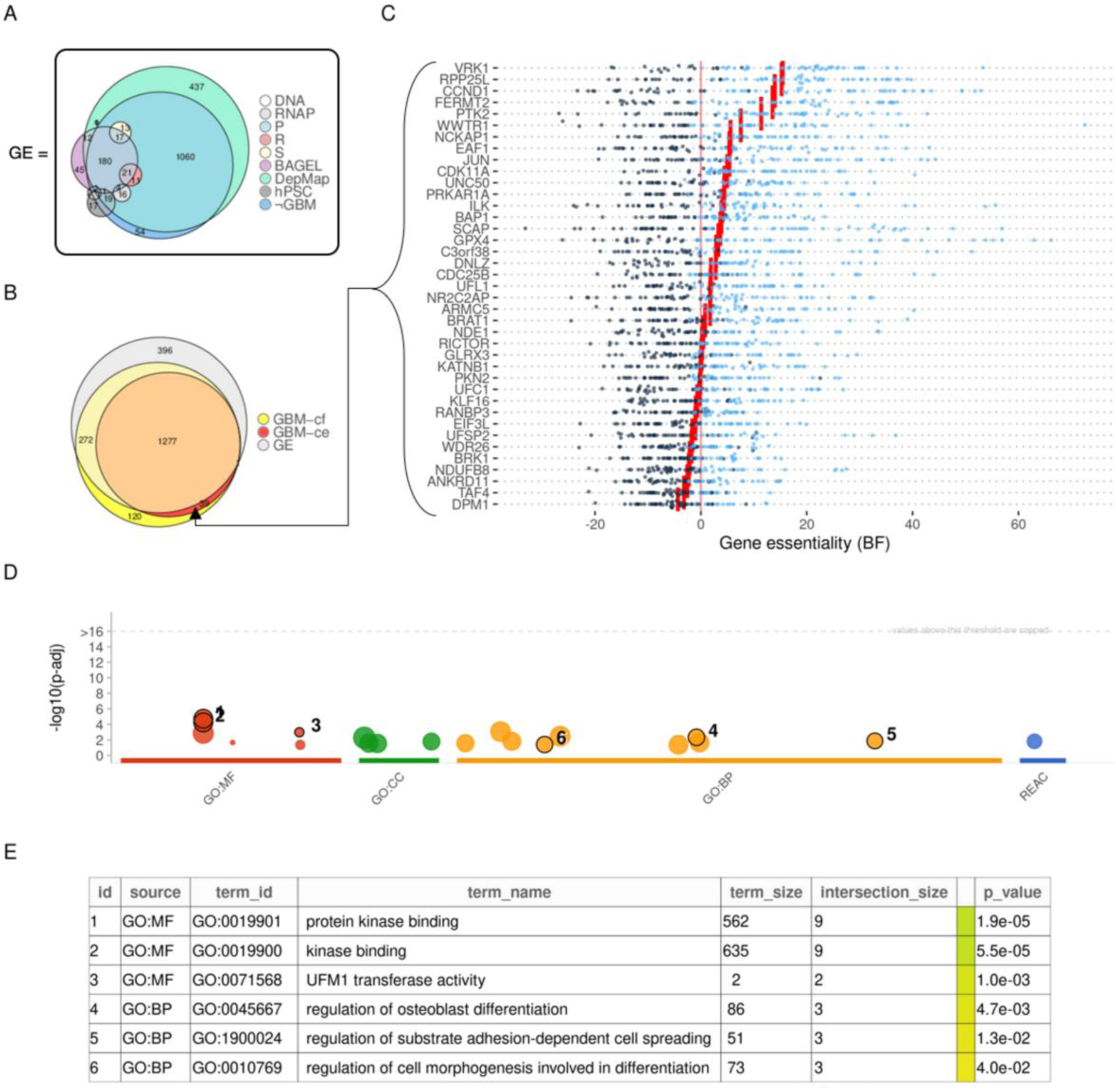
We curated a set of ‘generally essential’ genes to then define ‘GBM-specific commonly essential’ genes. [a] ‘Generally Essential’ (GE) genes were defined as the union of the following curated gene sets: DNA replication (DNA)^29^; RNA Polymerase (R)^29^; Proteasome^29^l; Ribosome (S)^29^; Splicosome (S)^29^; the CEGv2^90^ and the ‘curated’ BAGEL sets (BAGEL)^29^; the various essential genes defined within the DepMap (Common Essentials, Achilles Common Essentials and CRISPR Common Essentials) DepMap); genes that were essential in human pluripotent stem cells (hPSC)^28^; the results of our analysis to define common-essentials, and core-fitness genes in pan-cancer dataset excluding GBM and Glioma cell lines (¬GBM) (See Methods). [b] We determined GBM-core fitness (GBM-cf) and GBM-common-essential genes (GBM-ce) using Adam and FiPer^29^ respectively. We identified 39 GBM-specific commonly essential genes, defined as the set difference between the union of these gene sets and the GE genes (A) (See Methods). [c] Shows the essentiality scores for each GBM-specific commonly essential gene. Each dot represents a cell line; light blue dots indicate that the gene met the threshold for being essential in that cell line, whereas dark blue indicates that it did not. Red lines represent the median essentiality for that gene. [d] Enrichment plot for the GBM-specific commonly essential genes, highlighting the terms described in panel [e]. The most significantly enriched gene set was protein kinase binding.

### Annotating cell lines with candidate genetic and transcriptomic biomarkers that are relevant to GBM

GBM inter-and intra-tumoural heterogeneity has been studied extensively but remains incompletely understood. We used genomic and transcriptomic sequencing data to annotate each GBM cell line according to the presence or absence of common driver events including tumour-suppressor gene loss (loss-of-function mutation, copy-number deletion), oncogenic functional gains (oncogenic mutation, copy-number amplification)^11,12^, Telomerase Reverse Transcriptase (TERT) disinhibition^19^ and O-6-methylguanine-DNA methyltransferase (MGMT) suppression^18,30^ (see Methods). We used a modified single sample gene set enrichment algorithm (ssGSEA)^14^ to annotate each GBM cell line for the enrichment of various described GBM tumour-subtype signatures^13^ and GBM cell-state signatures^5,15–17^. We excluded cell lines with a mutation in IDH1 or IDH2 (IDH^mut^), in line with the modern consensus on the importance of IDH mutational status in GBM classification^9,10^. Because cancer cell lines^31^, and GBM cell lines in particular^32^ are prone to genetic drift over time, which can limit their clinical relevance, we also excluded any cell lines that were not enriched for at least one transcriptomic signature identified in GBM tumour samples^5,13,15–17^. Paralogous genes share sequence homology and functionality, and in some cases demonstrate mutual interdependency^33–37^. We identified genes which both (1) have known paralogues, and (2) are known to have low expression in some GBM contexts, as candidates for a paralogue-lethality relationship in GBM (see Methods). These lists comprised genes that are down-regulated as part of a certain GBM subtype^13^ or cell-state^5,15–17^, genes that are commonly methylated in GBM^20,21^, and genes on chromosome 10. We included the latter criteria because loss-of-heterozygosity (LOH) of chromosome 10 is a frequent driver event in GBM^22^ which leads to the passenger deletion of many genes. We annotated each GBM cell line based on the dichotomised expression of these genes (see Methods). Combining all of these analyses, we constructed a dataset detailing the presence of GBM-relevant biomarkers in GBM cell lines that had been subjected to genome-wide CRISPR/Cas9-mediated knockout screening (Fig 2).

**Fig 2:**
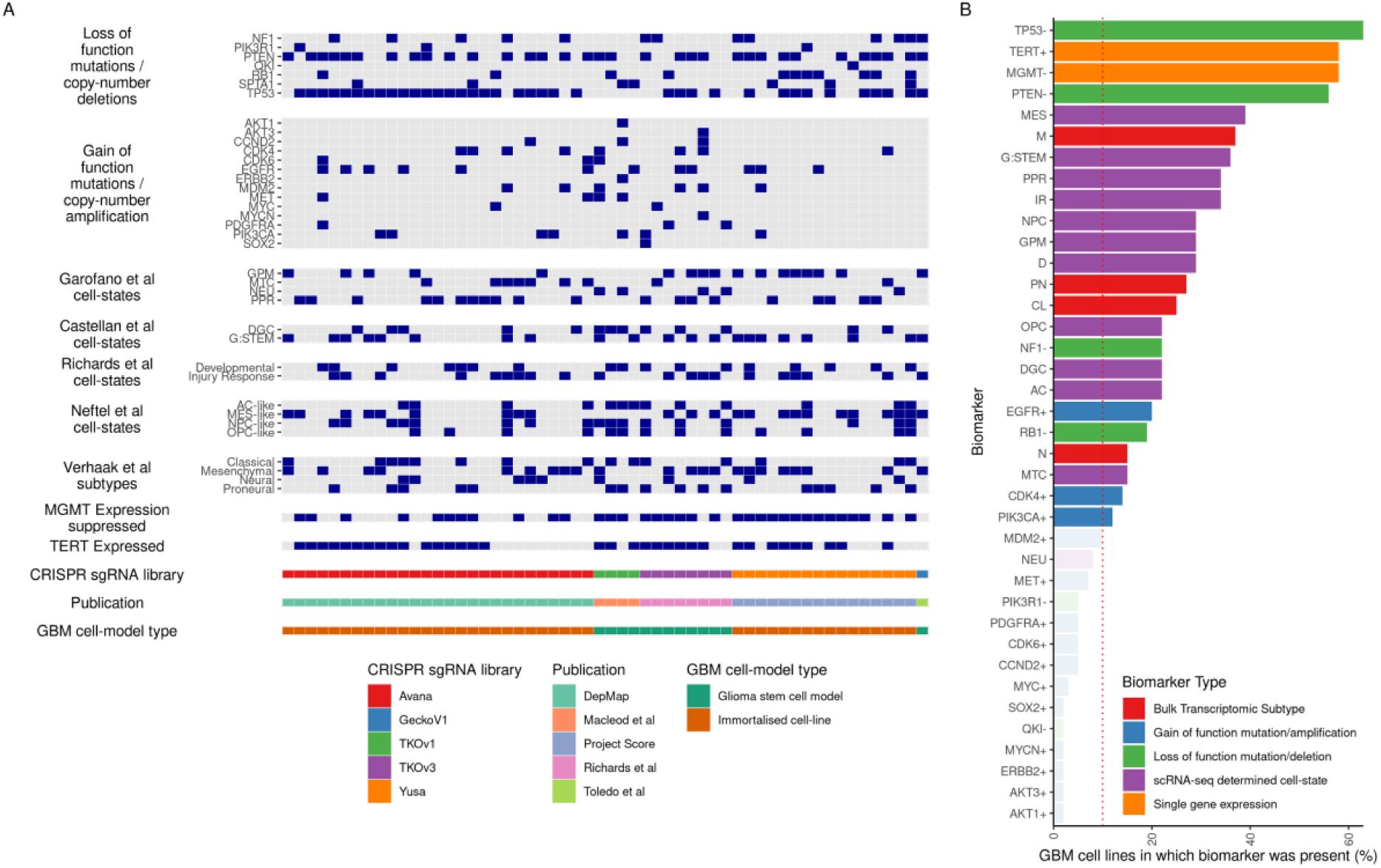
Curated dataset of biomarkers used in Targeted Learning analyses. [a] Graphical display of the curated biomarkers dataset used in our targeted learning analyses. We used WXS or WGS variant calls alongside RNA-seq data to infer the presence or absence of common genetic-driver events in GBM. We used ssGSEA to infer the presence or absence of numerous reported cell states identified through single-cell data analyses, or tumour subtypes identified through bulk-RNA sequencing analysis. We use RNA-seq data to infer MGMT-suppression status. We included potential batch effect confounders within the analysis including genome-wide pooled sgRNA library, study-of-origin, and GBM cell-model type. [b] Biomarkers were not evenly distributed amongst cell lines, but approximately replicated the distribution of prevalence in the human population. For example, TP53-, PTEN-, MGMT-suppression and TERT-expression are the most prevalent. However, all of the curated biomarkers shown are included as candidate confounder features during the SL training. We performed Targeted-Learning to determine the ATE and TSM only for biomarkers that were present in more than 10% of cell lines.

### GBM-CoDE uses Targeted Learning to robustly identify target-biomarker associations

Our objective was to robustly estimate the effect size for the influence of GBM-relevant biomarkers on gene essentiality, accounting for experimental heterogeneity, batch effects, and known biological interactions between biomarkers. For this, we employed Targeted Learning, a subfield of mathematical statistics and machine learning (ML), which utilises a model-agnostic ensemble ML framework, the Super Learner (SL)^38^. In general terms, the SL uses a diverse library of parametric and non-parametric ML algorithms to obtain an initial data-driven fit of the outcome of interest given the input data, whilst guarding against overfitting through cross-validation. This is followed by Targeted Maximum Likelihood Estimation (TMLE)^23^ which updates the initial SL fit, targeting it towards a specific quantity of interest, adjusting for confounders, to remove any residual bias due to model-misspecification^39^.

We performed Targeted Learning estimation procedures in a prioritized set of 536 genes we found to be highly essential (BF ≥ 6^26^) in GBM, but that were not ‘generally essential.’ To avoid selective inference^40^ in our subsequent Targeted Learning, we identified these genes using 14 partitioned GBM cell lines that had passed quality control but were ineligible for Targeted Learning either because they were duplicated cell lines between studies or had incomplete genetic or transcriptomic annotation (Fig 3a and b; see Methods). Our SL details the known biological interactions between genetic and transcriptomic variants that have been described in GBM, which the subsequent ML algorithms can utilise as features within their model (Fig 3c; see Methods). Following this initial step, one of two types of feature-screener algorithm subsets covariates based on their variable importance, before passing them to a library of candidate learning algorithms Fig 3d which are trained with up to 20-fold stratified cross-validation (Fig 3e; see Methods)^41^. We used this SL and TMLE to estimate two quantities of interest: the ‘average treatment effect’ (*ATE*), and the ‘treatment-specific mean’ (*TSM*) (Fig 3f; see Methods Equations (6) and (7) respectively). Here, the term *ATE* quantifies the effect size of a GBM-relevant biomarker on gene essentiality (quantified as a BF), adjusting for confounders, with confidence intervals. A positive *ATE* quantity indicates that when the biomarker was present in cell lines, the gene tended to be more essential than when the biomarker was absent. A negative *ATE* quantity indicates the inverse of this. Here, the term *TSM* quantifies the average essentiality of a gene (measured as a BF) in GBM cell lines when a biomarker is present adjusting for confounders.

**Fig 3:**
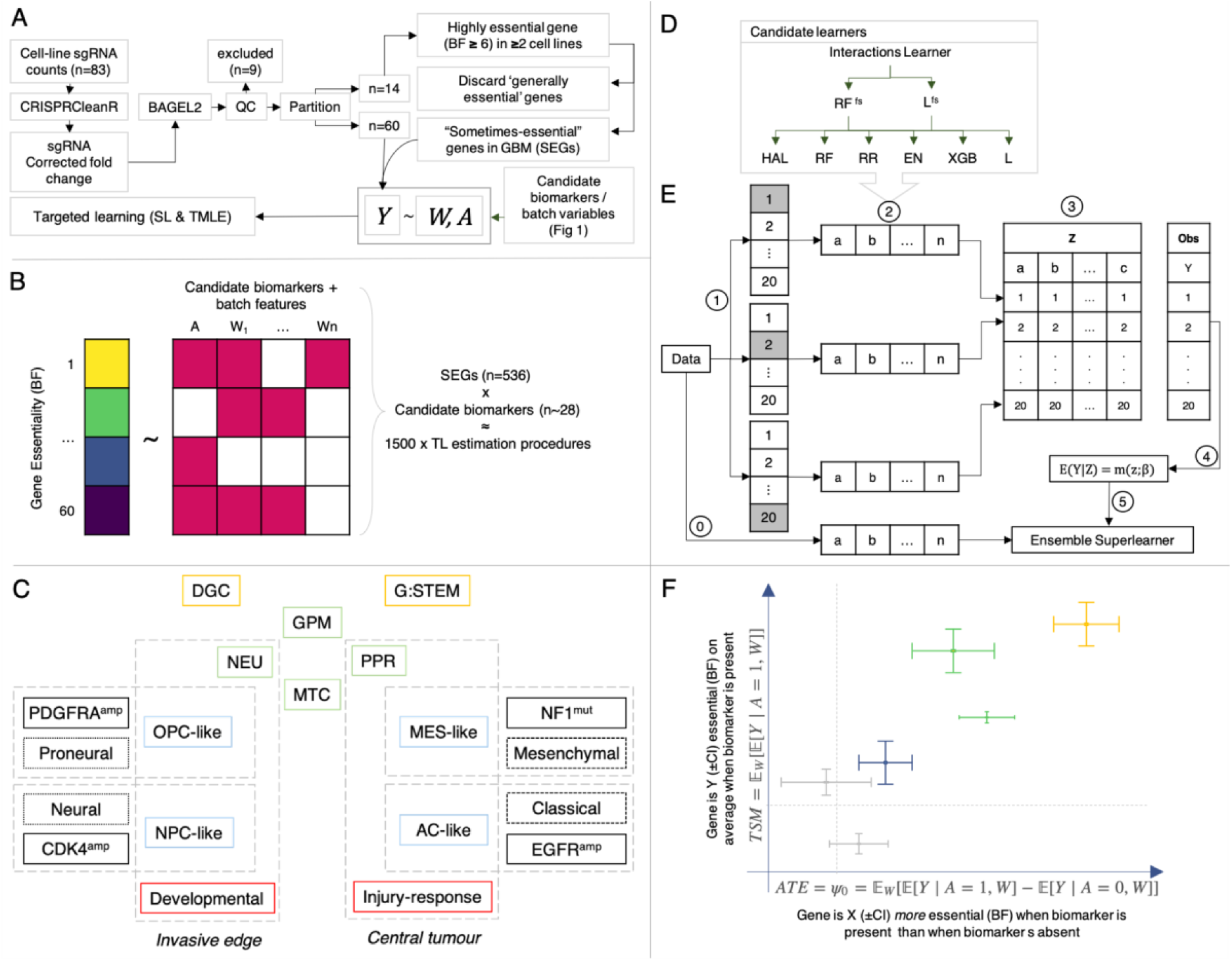
Summary of gene prioritisation and Targeted Learning algorithms. [a] We used BAGEL2^84^ to calculate essentiality (BF) using the CRISPRCleanR^24^ corrected sgRNA counts for each cell line. We partitioned 14 cell lines with incomplete genetic annotation and/or that were duplicated between datasets. We identified genes that were highly-essential (BF ≥ 6) in at least two of these partitioned cell lines. We excluded generally essential genes (See Methods). [b] With the remaining 536 prioritised genes, we used Targeted Learning to estimate the average treatment effect *ATE* of each biomarker (*A*) Fig 2 on gene essentiality (*Y*), adjusting for confounders (*W*). The data structure for each estimation is similar to regression of gene essentiality on covariates comprising curated biomarkers and batch features. [c] We supplied the candidate learners, with known biological interactions between known driver mutations (solid black) and transcriptomic classification systems: various described cell states (colour coded in yellow^16^, red^5^, green^17^, and blue^15^), and subtypes^13^ (dotted black). We also included interactions between batch effect features such as study, cell line, and sgRNA library (not shown). [d] The interactions are supplied to either a random forest (RF^fs^) or a lasso feature screener (L^fs^) followed by each candidate learner within our base learner library: Highly adaptive lasso (HAL), RF, Ridge regression (RR), Elastic net (EN) eXtreme Gradient Boosting (XGB) and Lasso (L). [e] The framework of a super (machine) learner (modified from)^110^ is as follows (0-5): 0: Train each candidate learner on entire dataset. 1: Split data into 20 folds for cross-validation stratified based on the biomarker *A*, or leave-one-out cross-validation for rarer biomarkers. 2: Train each candidate learner on every fold. 3: Predict the outcomes in the validation fold using the candidate learner trained on the testing fold. 4: Model selection and fitting for the regression of the observed outcome onto the predicted outcomes from the candidate learners. 5: Evaluate an ensemble SuperLearner by combining predictions from each candidate learner (step 0) with m(z;β) (steps 1-4). [f] The fits from each SL are used in cross-validated TMLE (CV-TMLE) to estimate both the *TSM* of biomarker presence, and *ATE* of the biomarker presence (*A* = 1) versus biomarker absence (A=0), adjusting for confounders (*W*) with confidence intervals.

Using this strategy we arrived at multiple target-biomarker pair hypotheses in GBM (Figs 5 and 4). Ultimately, the predictions from GBM-CoDE should be subject to extensive experimental validation and mechanistic study. However, fortuitously, two of the target-biomarker pair predictions that we identified have recently been experimentally validated by other groups in GBM^42–44^ and one has been mechanistically explained, albeit not in GBM specifically^45^ (see below; Fig 6). This implies that the additional, novel and currently unvalidated, findings from GBM-CoDE may be useful.

**Fig 4:**
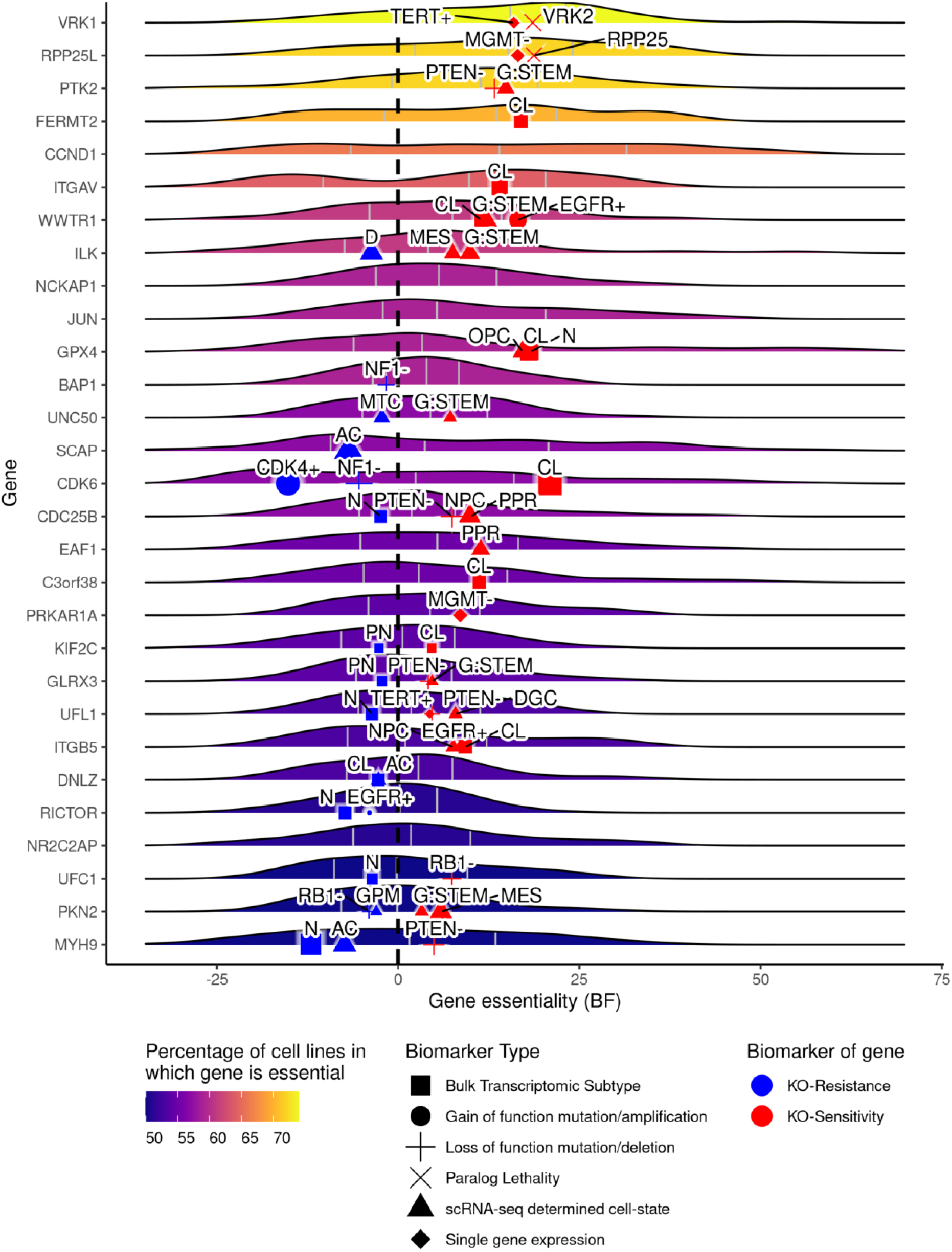
The distribution of essentiality scores for the most frequently essential genes in GBM, overlaid with the. *TSM* **for the biomarkers of gene knockout sensitivity and resistance.** Each ridge shows the distribution of essentiality scores (BF) for a gene across 60 GBM cell lines, with the median, first quartile and second quartiles overlaid. We show only genes that were essential in at least 50% of GBM cell lines and exclude ‘generally essential’ genes (see Methods). An adjusted BF of zero on the x-axis approximately corresponds to the minimum threshold for classifying a gene as essential; values greater than this mean that a gene was essential in a given cell line. Genes are ranked based on the proportion of cell lines in which they were essential, with VRK1 being the most commonly GBM-specific essential gene. Overlaid is the *TSM* for the most important biomarkers for each gene shown in either blue or red. Blue points (below 0 on the x-axis) are biomarkers of resistance to gene knockout and red points (above 0) are biomarkers of sensitivity to gene knockout. We determined the most important biomarkers for each gene based on the magnitude of the *ATE* on gene essentiality, (where it was statistically significant after the Benjamini-Hochberg Procedure) and the biomarker was associated with either a positive or negative *TSM* (indicating a binary classification of an essential or a non-essential gene respectively). Confidence intervals are not shown here for clarity but can be seen in Fig 5 The shape of each point corresponds to the type of biomarker. The label associated with each point details either the name of a specific gene or the name of a transcriptomic signature: N = Neural; CL = Classical; PN = Proneural; N = Neural; OPC = Oligodendrocyte precursor like^15^; NPC = Neural progenitor like^15^; MES = Mesenchymal like^15^; AC = Astrocyte like^15^; IR = Injury-response^5^; D = Developmental^5^; DGC = differentiated glioblastoma cells^16^; G:STEM = GSC-like^16^; GPM = glycolytic/plurimetabolic^17^; MTC = mitochondrial^17^; PPR = proliferative/progenitor)^17^.

**Fig 5:**
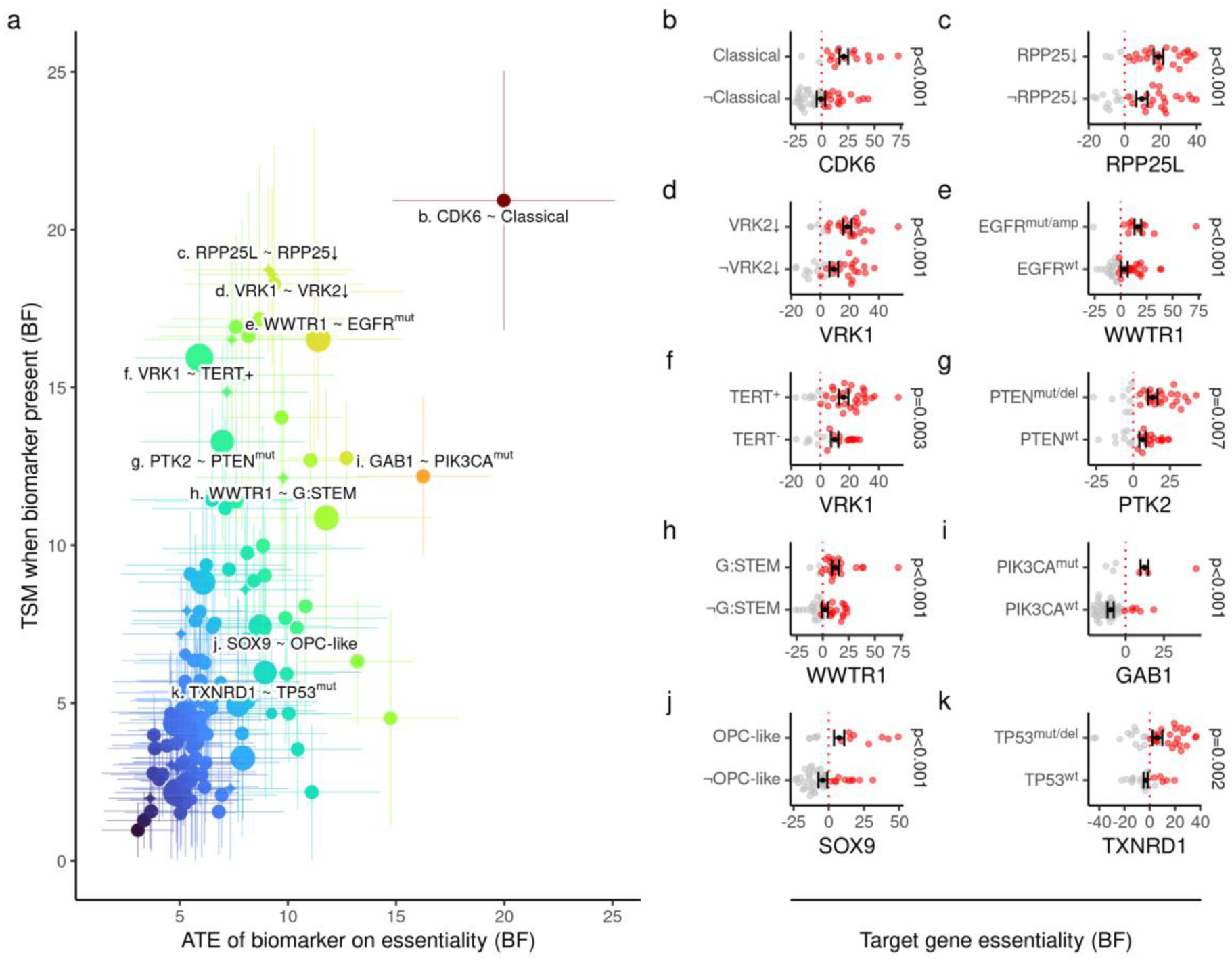
Identifying biomarkers which have the greatest magnitude association with increased gene essentiality. [a] Each point indicates a target-biomarker (TB) pair, some of which are labelled in the format ‘Target ∼ Biomarker’. The x-and y-axes represent the *ATE* and *TSM* of a TB pair respectively, with 95% confidence intervals (CI). TB pairs are coloured by ascending importance, quantified as the product of the *ATE* and *TSM*. The x-axis, can be thought of as follows: The essentiality of a given gene target (T), measured as a BF, was *ATE* ± CI higher in cell lines which contained a specific biomarker (B), than in cell lines which did not contain that biomarker. The y-axis quantity, can be thought of as follows: The essentiality of a given gene (T) was *TSM* ± CI in cell lines which contained a specific biomarker (B) than in cell lines which did not contain that biomarker. For a TB pair to be potentially therapeutically useful both the *ATE* and *TSM* must be positive. We label some TB pairs which are discussed in more detail within this manuscript. Within each of the panels b-h, the *TSM* ± CI of the gene essentiality in the presence and absence of the appropriate biomarker is shown (black), as is the with the Benjamini-Hochberg corrected p-values for the *ATE*. Additionally, in these panels, each dot represents a single cell line with its corresponding biomarker status (y-axis) and essentiality score (BF; x-axis), coloured by whether the gene was dichotomised as essential (red) or non-essential (grey). Two predicted TB pairs have recently been validated in GBM: [d] low VRK2 expression as a biomarker for VRK1 essentiality^42,43^ (see also Fig 6a-c), and [e] EGFR^amp/mut^ status as a biomarker for WWTR1 essentiality^44^ (see also Fig 6g). Additionally, the paralogue lethality relationship between RPP25L and RPP25 [b] has been mechanistically validated^45^ albeit not in GBM specifically (see also Fig 6d-f).

**Fig 6:**
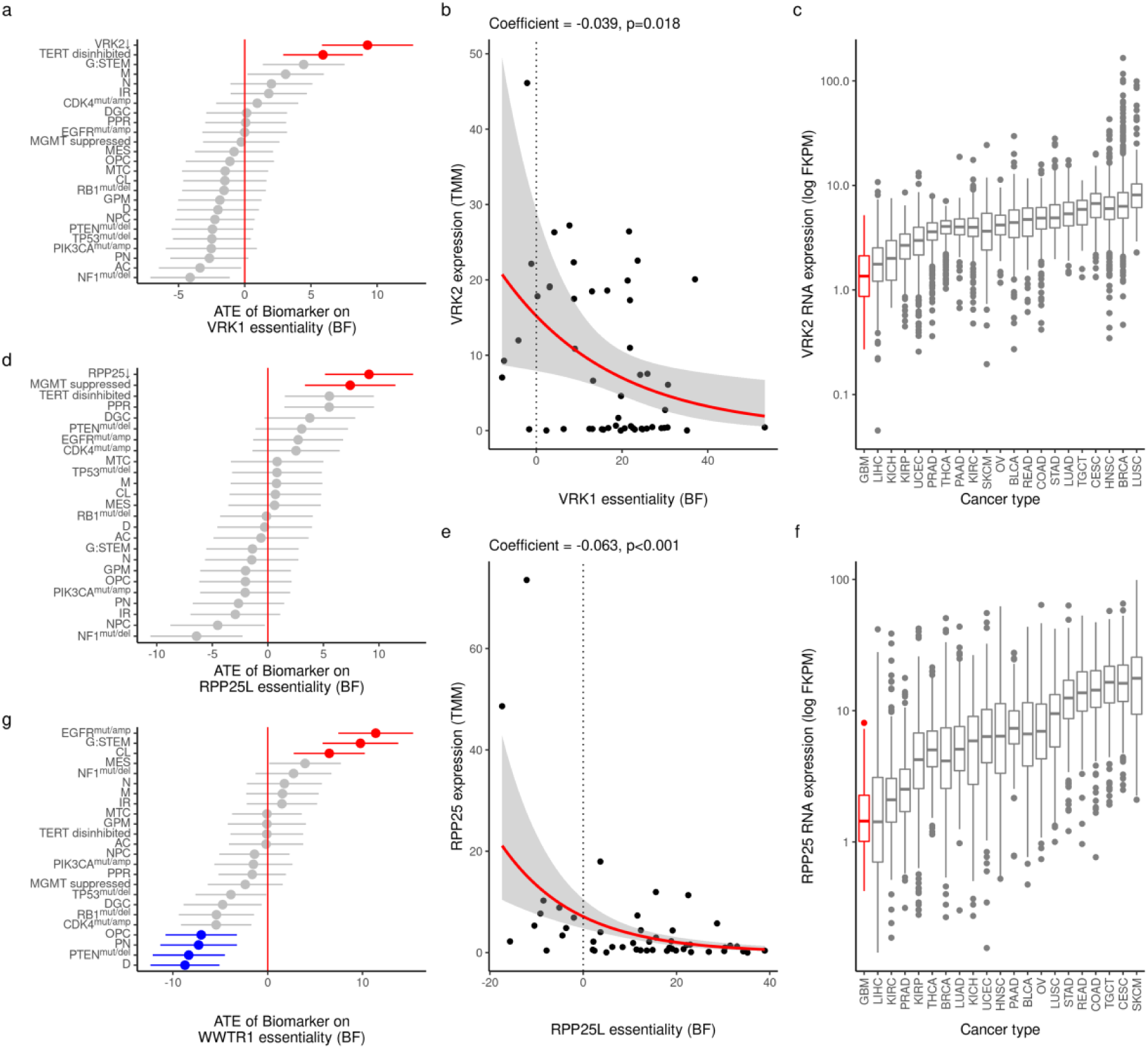
Three target-biomarker pairs have orthogonal mechanistic validation, two specifically in GBM. [a] The biomarker that was most associated with increased VRK1 essentiality, was the low expression of its paralogue VRK2. This paralogue-lethality relationship has been validated in GBM^42,43^. [b] Using a negative binomial model, VRK1 essentiality was inversely associated with VRK2 expression. [c] Of all cancer types in the TCGA dataset, GBM had the lowest expression of VRK2. [d] The biomarker which most increased RPP25L essentiality, was the low expression of its paralogue RPP25. The mechanism behind this paralog-lethality relationship has been described^45^ albeit not in GBM specifically. [e] Using a negative binomial model, RPP25L essentiality was inversely associated with RPP25 expression. [f] Of all cancer types in the TCGA dataset, GBM had the lowest expression of RPP25. [g] WWTR1 was more essential in cell lines with EGFR^mut/amp^ status. This target-biomarker relationship has been validated in GBM^44^, leading to a biomarker-stratified randomised controlled trial^52^. Additionally, we identified the G:STEM cell state^16^ was associated with increased WWTR1 essentiality. Mechanistically, this is likely due to the role of WWTR1 as a master regulator of the G:STEM cell state^16^. Abbreviations: N = Neural; CL = Classical; PN = Proneural; N = Neural; OPC = Oligodendrocyte precursor like^15^; NPC = Neural progenitor like^15^; MES = Mesenchymal like^15^; AC = Astrocyte like^15^; IR = Injury-response^5^; D = Developmental^5^; DGC = differentiated glioblastoma cells^16^; G:STEM = GSC-like^16^; GPM = glycolytic/plurimetabolic^17^; MTC = mitochondrial^17^; PPR = proliferative/progenitor)^17^.

### GBM-CoDE successfully discovers independently validated target-biomarker pairs

VRK1, which encodes VRK Serine/Threonine Kinase 1, was essential in 86% of GBM cell lines, making it the most frequently essential ‘GBM-specific commonly-essential’ gene Fig 4. VRK1 is a chromatin kinase that regulates DNA damage responses, regulates chromatin remodelling^46^, and is a downstream target of SOX2 regulation^47^, the latter being important in Glioma stem cell identity regulation^48^. The primary biomarker we identified for VRK1 essentiality, was low-expression of its paralogue VRK2 Fig 6A. Accordingly, there is an inverse correlation between VRK1 essentiality and VRK2 expression Fig 6B. VRK1 dependence may be especially useful in GBM, where VRK2 is commonly hypermethylated^20^ and has lower expression than in other cancers Fig 6C. This paralogue-lethality relationship has been identified and validated in GBM by two groups independently^42,43^. Further research is warranted to establish whether, and if so how, low-expression of VRK2 affords a fitness advantage for a GBM cell. Additionally, we identified TERT disinhibition as a biomarker of increased VRK1 essentiality Fig 6A. Disinhibition of telomerase activity, most commonly due to TERT-promotor mutation, is a common event in GBM^49^. While this target-biomarker pair has not been described in GBM, and further experimental validation is required, it is mechanistically plausible given that VRK1 deficiency induces telomere shortening^50^.

WWTR1 (also known as TAZ), which encodes WW domain-containing transcription regulator protein 1, was essential in 70% of GBM cell lines. WWRT1-knockout was more essential in GBM cell lines when an EGFRvIII-variant mutation^51^ (EGFR^vIII^) or EGFR-amplification (EGFR^amp^) - EGF^mut/amp^ - was present, than in EGFR-wildtype (EGFR^wt^) cell lines Fig 6G and H. This target-biomarker pair is potentially important because EGFR^mut/amp^ is found in 57% of GBM samples^11^. One of the few recruiting clinical trials in GBM that is biomarker-stratified is testing Verteporfin (which inhibits WWTR1 transcriptional activity) in EGFR^mut/amp^ GBM^52^. This trial is supported by preclinical data demonstrating the vulnerability of EGFR^vIII^-containing GBM cell lines to short-hairpin-RNA mediated knock-down of WWTR1^44^. As an additional level of validation, we identified the ‘G:STEM’ cell state^16^ was a biomarker of WWTR1 essentiality Fig 6G. Mechanistically, this is likely related to the role of WWTR1 as a master regulator of the G:STEM cell state^16^.

RPP25L, which encodes Ribonuclease P/MRP Subunit P25-Like protein, was essential in 82% of GBM cell lines Fig 4. The biomarker that had the largest magnitude effect on RPP25L essentiality, was low-expression of its paralogue gene RPP25, which encodes Ribonuclease P And MRP Subunit P2 Fig 6D. Accordingly, RPP25L essentiality is inversely correlated with RPP25 expression Fig 6E. We note that RPP25L can physically and functionally compensate for RPP25 within the RNase P/MRP complexes that regulate tRNA processing^45^, explaining the mechanism behind this paralogue-lethality. This may be exploitable in GBM, in which RPP25 is commonly methylated^21^, and has lower expression than other cancers Fig 6F.

### GBM-CoDE identified putative context-dependent vulnerabilities targetable with existing drugs

We highlight three target-biomarker pair results Fig 5d, e and g that may be rapidly clinically translatable if experimentally validated; hypotheses in which the biomarker was a common genetic driver in GBM, the gene product of the target can be inhibited with existing small molecule drugs, and there is a plausible mechanism behind the putative target-biomarker relationship based on existing biological knowledge: (1) Target: PTK2, Biomarker: PTEN^mut^ Supplementary Fig S1 (2) Target: TXNRD1, Biomarker: TP53^mut^ Supplementary Fig S2, and (3) Target: GAB1, Biomarker: PIK3CA^mut^ Supplementary Fig S3. Common driver mutations may be important biomarkers because, while a GBM cell may transition between various cell states in response to stress, it cannot easily evolve to re-instate the function of tumour suppressor gene once it has been mutated or deleted, nor can it easily relinquish an oncogene addiction.

CDK6, which encodes Cyclin-dependent kinase 6, was essential in 62% of GBM cell lines. It had very high essentiality scores in cell lines that were enriched for the ‘Classical’ GBM subtype^13^ (Fig 5a; Fig 8a). There was a positive correlation between CDK6 essentiality and multiple genes that form the Classical subtype signature^13,14^ Fig 8e-i. We identified CDK4- amplification (CDK4^amp^) as the strongest biomarker of resistance to CDK6-knockout Fig 8A. Mechanistically, this is likely due to the paralogous relationship between CDK4 and CDK6^53^. If experimentally validated, this may suggest CDK6 inhibition could be useful in patients who have tumours that are enriched for the ‘Classical’ subtype, but that do not have a CDK4^amp^. However, we note that clinically approved CDK6 inhibitors also inhibit CDK4, which may limit their therapeutic window due to paralogue-lethality-associated toxicity to normal tissues.

**Fig 7:**
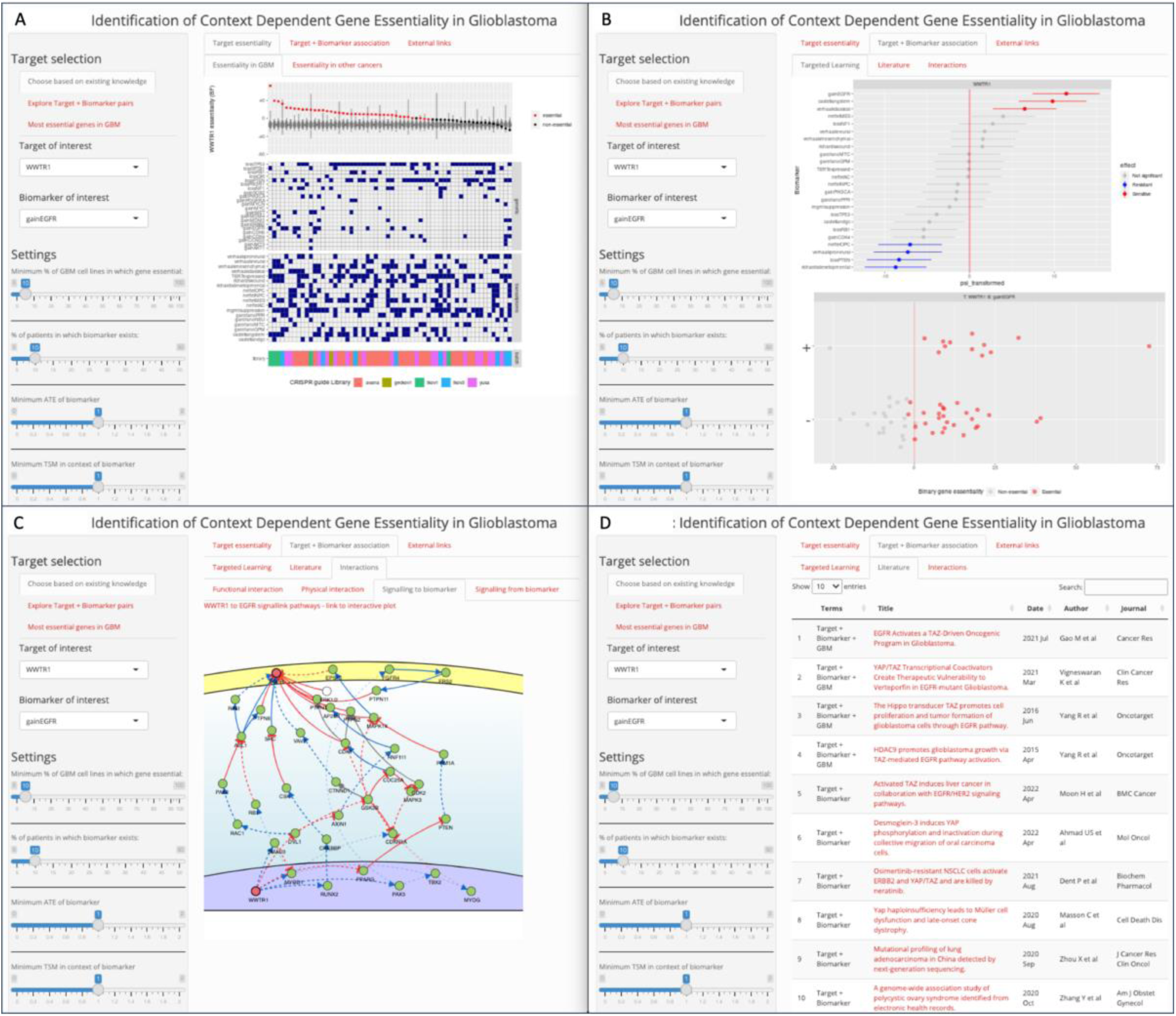
The results from GBM-CoDE can be explored interactively to prioritise T:B pairs for validation. An example exploration of this data in the GBM-CoDE app, looking at WWTR1 as the target and EGFR^mut/amp^ as the biomarker: gene level essentiality alongside the cell line’s genetic and transcriptomic features [a], statistical modelling results [b], Known signalling links between WWTR1 and EGFR [c], and existing literature linking WWTR1 and EGFR ± GBM [d].

**Fig 8:**
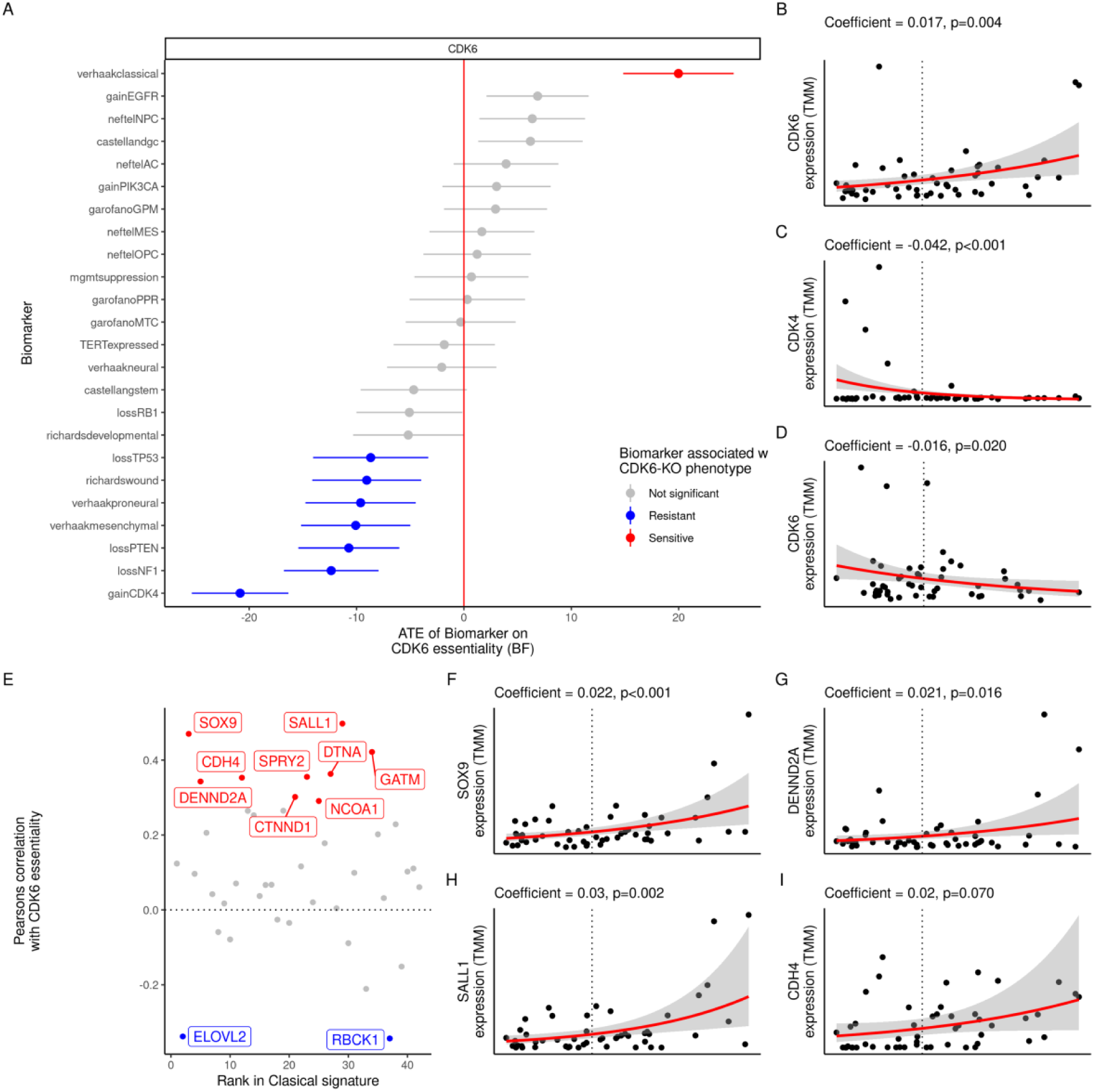
CDK6 knockout sensitivity and resistance were associated with the Classical subtype, and CDK4^amp^ respectively. [a] Targeted-Learning identified the ‘Classical’ subtype transcriptomic signature^13^ as having the largest *ATE* on CDK6 essentiality. Additionally, we identified that CDK4 amplification was associated with resistance to CDK6-knockout tolerance. [b]: Using negative binomial regression, we identified a positive correlation between CDK6 essentiality and CDK6 expression. [c]. CDK6 essentiality was inversely correlated with CDK4 expression, and visa versa [d]. [e]: We found that the expression of multiple genes within the Classical-subtype signature^14^ showed a positive correlation with CDK6 expression. We confirmed this using negative binomial regression; shown here are the scatter plots with associated coefficients and p-values for SOX9 (F), DENND2A (G), SALL1 (H) and CDH4 (I).

### SOX9 is more essential in the context of the cell states that predominate at the invasive edge of GBM

Heterogeneous cells within a single GBM tumour have been described as resembling various neurodevelopmental cell types^15^. Some of these cell states, including the OPC-like and NPC-like cell states, are found more frequently within the invasive margin of GBM than in the tumour core^54^. We used GBM-CoDE to probe for prominent dependencies associated with these cell states Fig 9. The gene in which the OPC-like cell state had the largest *ATE* on essentiality, was SOX9 (Fig 5a and f, and Fig 9a). SOX9 encodes SRY-box transcription factor 9, which is involved in neurodevelopment, and has an important role in Glioma stem cell maintenance^55^. Additionally, the NPC-like cell state was associated with increased SOX9 essentiality Fig 9b. We note that CRISPR-Cas9-mediated-knockout of SOX9 was not deleterious in human neural stem cells^56^, or human pluripotent stem cells^28^, nor was CRISPR/dead-Cas9-mediated repression (CRISPR-interference; CRISPRi) of SOX9 transcription in normal glutamatergic neurons^57,58^. Together, these findings suggest that therapeutically inhibiting the function of SOX9 may perturb GBM cells that adopt an OPC-like or NPC-like cell state, without inducing neural or neural stem cell toxicity. It follows that doing so in human GBM patients may perturb post-operative residual cells within the invasive edge, reducing their capacity to drive recurrence from the resection margin. However, extensive experimental validation is required, and we note that it is difficult to inhibit transcription factors therapeutically.

**Fig 9:**
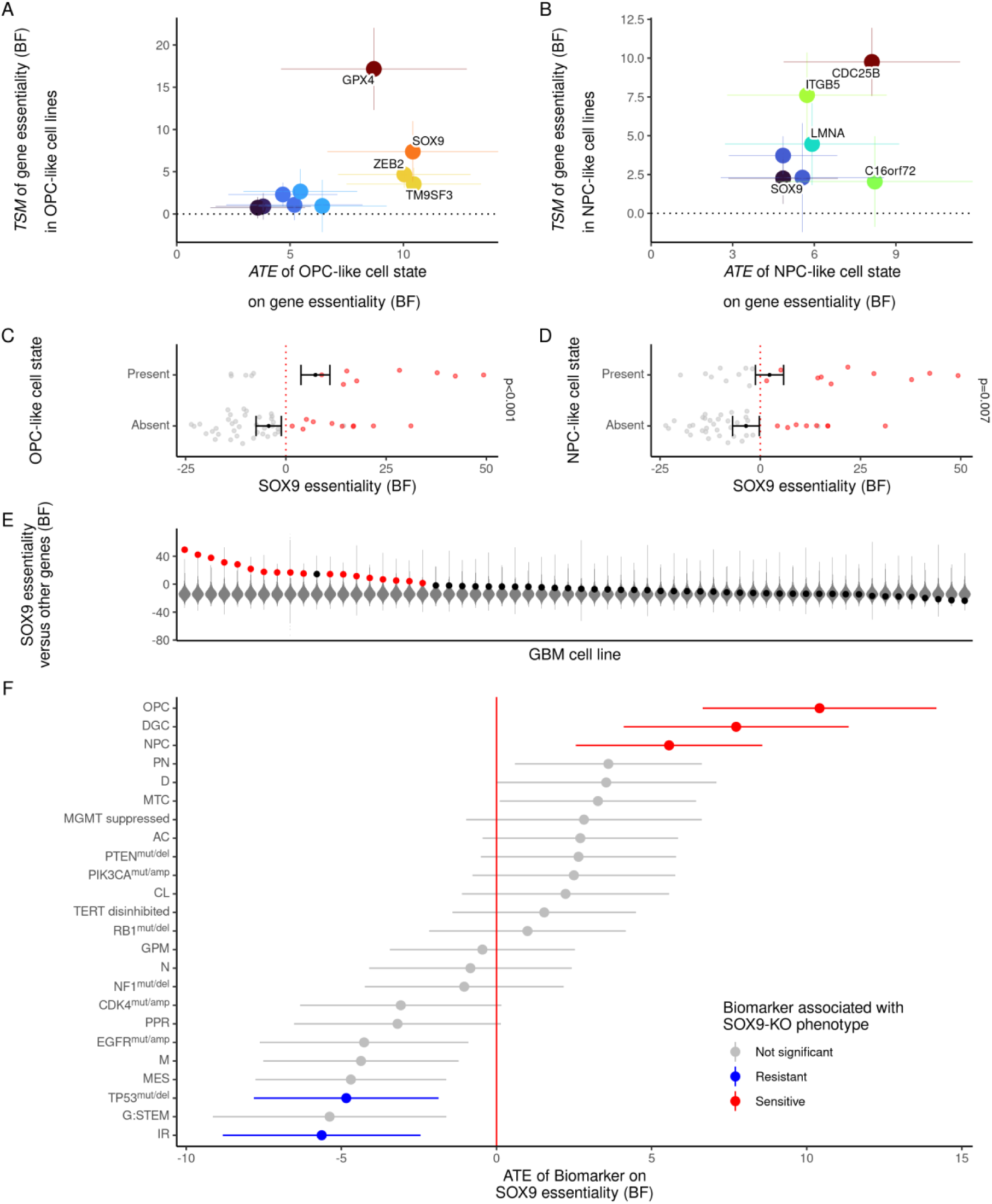
SOX9 is more essential in the context of the OPC-like and NPC-like cell states; states which predominate in the invasive edge of GBM. We wanted to identify which genes were most influenced by cell states that have been reported to predominate in the invasive edge of GBM, including the OPC-like cell state and NPC-like^54^. [a] We identified genes in which the OPC-like cell state had the largest positive *ATE*. [b] We identified genes in which the NPC-like cell state had the largest positive *ATE*. [c] SOX9 essentiality was higher in cell lines that enriched for the OPC-like cell state, as well as in cell lines that enriched for the NPC-like cell state (d). [e] SOX9, was essential in only 43% of GBM cell lines, and was not identified to be commonly essential in GBM. [f] Forest plot indicating the ranked *ATE* of each biomarker on SOX9 essentiality. GBM cells are reported to exist on an axis between ‘proneural’ and ‘mesenchymal’ states^111^ We note that transcriptomic signatures which may be considered within a broader ‘proneural’ group (including OPC-like, NPC-like, Proneural, Developmental) tended be associated with higher SOX9 essentiality, and those which may be considered within a broader ‘mesenchymal’ group (including Injury-response, MES-like, Mesenchymal, and G:STEM) tended to be associated with lower SOX9 essentiality, albeit not all of these were statistically significant after the Benjamini-Hochberg Procedure. These putative target biomarker pairs are potentially important because they imply that SOX9 antagonism may perturb cells which adopt an NPC-like or OPC-like state, states are described to predominate in the invasive edge of GBM. However, Further experimental studies are required. Abbreviations as per Fig 6.

## Discussion

We developed GBM-CoDE, a target-biomarker discovery platform for GBM comprising curated datasets, interrogated using a Targeted Learning driven algorithm to identify context-dependent gene essentiality. Each of these target-biomarker pair hypotheses, if experimentally validated, could theoretically accelerate towards a clinical trial of the format: [target inhibitor] versus placebo in [biomarker-positive] GBM. To democratise the experimental validation of these putative target-biomarker pairs, we will release them freely in a graphical user interface (Fig 7a and b). There, researchers can visualise the known relationships^59,60^ between relevant biological entities which may give insight into the potential underlying mechanism behind the putative target-biomarker pair Fig 7c and d.

Our approach builds upon other publications that explore biomarkers of gene essentiality^4,27,33,33,61–63^. To our knowledge, our analysis is the first to deliberately compromise dataset size in exchange for specificity and quality, aiming to capture the heterogeneity of a complex disease as accurately as possible. Secondly, we acknowledge that driver mutations and cell states frequently co-occur^15^. Therefore, our approach incorporates features of known biological significance, as well as the known relationships between these, whilst providing mathematical guarantees on the resulting estimates. This allowed us to include batch variables as independent features in our statistical modelling rather than batch correcting whole datasets^27^, aiming to maximise the opportunity to identify context-dependent gene essentiality.

There are some limitations to our analysis. Firstly, the datasets that we integrated were heterogeneous and we did not have access to every form of raw data for every cell line. Secondly, our approach aims to identify cancer cell-intrinsic genetic vulnerabilities only. This is one of, but not the only challenge to developing successful therapies in GBM; we do not aim to address: the role of the tumour microenvironment (including immune evasion^64^, and neuronal hijacking^54,65^); the need for therapeutics to either penetrate the blood-brain-barrier; or the need to synergise with standard treatment.

Although we present some external validation of three target-biomarker pair hypotheses^42–45^ Fig 6, the majority of our hypotheses are novel, and remain unvalidated; and extensive experimental validation is required. We have built GBM-CoDE so that as more functional genomic data becomes available, and as more knowledge of GBM heterogeneity arises, we can include this in updated versions. Finally, while GBM is an ideal type of cancer upon which to use it, our method is broadly generalisable to other cancers.

## Methods

Supplementary Table 2 details key resources, software and data used in the preparation of this manuscript.

### Collation and analysis of genome-wide pooled CRISPR-Cas9-mediated knockout screens in GBM

#### Literature review to identify datasets containing CRISPR screens in GBM cell lines

We reviewed the literature for studies that performed genome-wide pooled sgRNA CRISPR/Cas9-mediated knockout screens in GBM cell lines, identifying six candidate papers^4–8,66^.

Details of the datasets are provided in Table 1, but briefly, they comprise the following. The “Depmap”^67^ is an international, multicentre collaborative project which involves performing drug sensitivity screens, and genome-wide loss-of-function screens on genetically annotated cell lines of multiple cancer types including GBM. We identified cell lines representative of GBM within the ‘AVANA’ sgRNA library CRISPR-KO screening dataset^8,68,69^. “Project Score” is a similar large-scale pan-cancer effort from the Sanger institute^4,70,71^ where multiple cancer cell lines including GBM were screened using the ‘Yusa’ library^72,73^. We identified other papers in which CRISPR-KO screening was performed specifically in glioma stem cell (GSC^74^) lines: Toledo et al^7^ (2 GSCs; ‘GeCKO’ library (Version 1.0)^75^, Macleod et al^6^ (8 GSCs; TKOv1^66,76^ & TKOv3^26,77^) and Richards et al^5^ (11 GSCs; TKOv3). We identified that one GSC line screened by Hart et al^66^ was also screened in Macleod et al^6^ and we, therefore, did not include this paper in our analysis. From these datasets, we identified cell lines which had been subjected to whole-genome CRISPR-KO screening experiments, which may be eligible for inclusion in our study.

#### Source data experimental methods: generation of Cas9-expressing GBM cell lines to subject to pooled whole-genome CRISPR sgRNA perturbation

Full details of the experimental methods used to derive Cas9-expressing cancer cell lines upon which to perform whole-genome CRISPR screen are available within the source data papers^4–8,66^. In the studies using GSC models^5–7^, established serum-free culturing methods were used to generate patient-derived GSCs^74^. GBM cell lines were engineered to express Cas9 nuclease, via transduction with a lentiviral vector expressing Cas9 nuclease, followed by selection for Cas9 expressing cells through antibiotic resistance detection, followed by a Cas9 activity assay (Table 1). Cas9 expressing cancer cell lines were then transduced with a lentiviral packaged whole-genome sgRNA library, and at the designated time endpoint, DNA was extracted, sgRNA PCR amplification performed, and sequencing performed (Table 1).

#### Computational methods

##### CRISPR-KO sgRNA read-count data processing and copy number correction

We collated and cleaned sgRNA read-count data for each cell line, and then we performed copy number correction using CRISPRCleanR (v2.0.0) using the default settings^24,78^.

To enable this we generated CRISPRCleanR libraries for the GeckoV1 TKOv1, TKOv3, AVANA and Yusa sgRNA libraries. We aligned sgRNA sequences to GRCh38^79^ with Bowtie (v1.3)^80^, then mapped the sgRNA perfect matches to target genes in GENCODE GRCh38.p13 (Release 35)^81^. We discarded sgRNAs that did not align perfectly to *any* genomic location. Where the targeted genes did not align with the original authors’ target gene, we checked for synonyms in updated gene symbols and updated where appropriate. Where the sgRNAs targeted multiple genes at the same location (most commonly at readthrough or overlapping genes) we chose the gene most commonly identified as the target by other guides.

All gene names within datasets (sgRNA libraries, CRISPRCleanrR libraries, BAGEL2 multi-target-creation libraries, and reference gene lists) were updated using HGNChelper^82,83^.

##### Calculating gene essentiality (BAGEL2)

With the copy-number-corrected counts, we generated a gene-level Bayes Factor of essentiality for each cell line using BAGEL2 with multi-target correction mode enabled^84,85^.

The Bayesian Analysis of Gene Essentiality (BAGEL) algorithm quantifies gene essentiality based on functional genomic experiments^84^. Using reference gene sets as Bayesian priors, BAGEL computes the likelihood of a gene’s membership to either an “essential” or the “non-essential” gene class^84^. It outputs a log Bayes Factor (BF) representing the union of biological effect size (in this case cell survival after perturbation) and statistical confidence^84^.

To enable this we created multi-target-correction libraries for GeckoV1 TKOv1, TKOv3, AVANA and Yusa sgRNA libraries using GENCODE GRCh38.p13 (Release 35)^81^, and the reference genome GRCh38^79^. and the ‘pre-calculation of library alignment info’ within BAGEL2^85,86^.

##### Reference essential gene sets

We noted that some of the reference ‘non-essential genes’ were essential (sometimes highly so) in numerous GBM cell lines. This observation was particularly evident in GSC lines. For example, OLIG2 is part of the reference “non-essential” gene set. It was’essential” (FDR<0.05 on BAGEL2 precision-recall) in 8 GBM cell lines (11% of those studied). OLIG2 is a CNS-specific transcription factor involved in glial and neural progenitor proliferation and specification^87^. OLIG2 is frequently expressed in gliomas, with an important role in gliomagenesis^87^. OLIG2 silencing in patient-derived models results in a shift from a proneural to classical or mesenchymal phenotype^87^. Given its CNS-restricted expression it follows that it would be non-essential in non-CNS tumours, but knowing the above, it follows that it may be *sometimes* essential in GBM, and therefore not *non*-essential.

Because of the relative under-representation of Glioblastoma in the datasets from which the ‘essential’^26^ and ‘non-essential’^88^ gene sets were derived, we were concerned that using them without curation may bias the Bayesian estimates of essentiality for genes of interest where essentiality status is unknown. To address this, we generated a curated set of ‘essential’ and ‘non-essential’ genes that were appropriate to Glioblastoma cellular models, in particular GSCs. To do this, we used the Adaptive Daisy Model (ADaM^4,89^), run with default settings. We ran ADaM with four iterations: Using the ‘Project-Score’ ‘pan-cancer core-fitness’ genes^4^ and the ‘Hart’ essential genes^66^ as reference essentials on the GSC-models only, and again on all Glioblastoma models that passed biological QC. We then defined ‘agreed essentials’ as the intersection of both the Hart reference essential genes^66^ and Sanger core-fitness genes^4^. An essential gene was retained in our curated reference set if it was 1: an ‘agreed essential’ gene, and 2: found with ADaM to be a core-fitness gene in the GBM cell lines studied. Additionally, we retained genes that were context-specific fitness genes in the GSCs from *either* the Hart or the Sanger reference essentials. Genes were discarded from the reference ‘non-essential’ gene set if they were found to have a FDR<0.05 and a BF of three or more in four or more cell lines that passed biological QC. Non-essential genes were also discarded if they constituted core-essential or common-essential genes in Project Score or the Depmap (20q3) respectively. Finally, we used these custom reference sets in determining which cell lines were of sufficient technical quality for inclusion. However, to avoid self-referentiality and selective inference, we did not use this in the main Targeted learning analysis, where we instead used BAGEL-curated gene sets^29^, in which OLIG2 is correctly removed from the non-essential gene list, among others.

##### CRISPR-KO screen technical quality control (Cohen’s D and F-measure)

One method for determining the technical quality of a replicate within a screen is to determine the degree of overlap between the distribution of the gene essentiality scores in the reference essential versus reference non-essential genes, which can be measured with the Cohen’s D score^84^ ((1)):

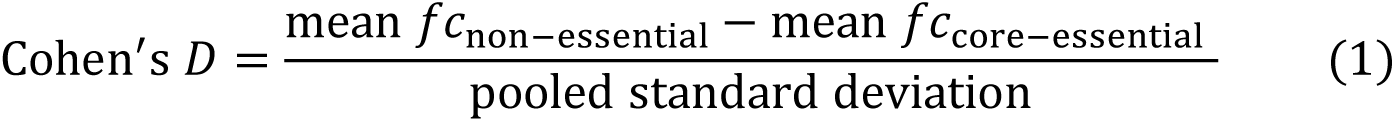

We used this method to determine if any technical replicates should be excluded from the analysis. We used the F-measure (the harmonic mean of precision and recall at the threshold of BF=5) as a metric of essentiality screen result quality^84^. Cell lines with an F-measure of less than 0.7 were excluded for failing technical QC. We excluded 5 cell lines from our analysis based on this technical QC metric.

##### Removal of duplicated cell lines for statistical modelling

Some cell lines were duplicated across studies. We anticipated this would cause bias during statistical analysis for context-dependent gene essentiality. The proportion and significance of associations between genetic events and gene essentiality would be artificially high in lines that were duplicated between studies. Therefore, where cell lines were duplicated in Richards et al from Macleod et al, we retained the Richards et al (TKOv3 CRISPR library) screen because the TKOv3 library is an improvement on the TKOv1 library. In lines that were duplicated across the DepMap and Project-Score datasets, we chose the screen with the highest mean Cohen-D score across replicates.

##### Integration of CRISPR-KO screen data into a binary matrix of gene-level essentiality

For each cell line, genes with an essentiality BF false discovery rate (FDR) of p ≤ 0.05 were labelled ‘essential’ and others ‘not-essential.’ FDR is quantified as FP/(FP + TP) with positives and negatives as per the reference gene sets^90^. This was used to compile a binary matrix of essentiality.

##### Integration of CRISPR-KO screen data into a matrix of gene-level essentiality

We used an established method to adjust BF values for ease of interpretation^4^. This involved finding the minimum BF where FDR was <0.05 on precision-recall and subtracting this from all BF values, such that in most circumstances all essential genes had a BF greater than 0 and non-essential genes had a BF less than or equal to 0. Genes that were not part of the screening library in all cell lines were discarded prior to integration. We integrated data from the cell lines that passed biological and technical QC, and the best candidates in the case of duplication as described. Bayes factors had been normalised on a per-screen basis from the BAGEL2 algorithm. We performed quantile normalisation to facilitate comparison between cell lines. We did not perform batch correction on this dataset, instead opting to use various batch features in our subsequent targeted learning.

### Identifying ‘GBM-Specific Commonly-Essential’ genes

We generated a list of ‘non-GBM core-fitness genes’ (¬*GBM_cf_*) and ‘non-GBM commonly-essential genes’ (¬*GBM_ce_*) using the Adaptive Daisy Model (ADaM^29^) and the Fitness Percentile AUC method (FiPer^29^) algorithms respectively, applied to a subset of an integrated pan-cancer dataset^27^, which excluded Glioma cell lines. This is because genes that are commonly-or core-essential in non-GBM cancers, but not in GBM, may be masked in existing pan-cancer reference essential lists that were created using datasets which included GBM cell lines. We excluded all Glioma cell lines (not just GBM cell lines specifically) because Glioma is a superset of brain tumours that includes GBM. Next, we identified existing lists of reference essential (*RE*) genes Equation (2): the CEGv2^90^ and the ‘curated’ BAGEL sets (BAGEL)^29^; genes involved in DNA replication, RNA Polymerase, Proteasome, Ribosome, and the Splicosome^29^. We defined ‘Generally Essential’ (*GE*) genes as the union of these reference essential genes (*RE*), various essential genes identified within the DepMap (Common Essentials, Achilles Common Essentials and CRISPR Common Essentials; *CE_DepMap_*), the human pluripotent stem-cell ‘essentialome’ (ℎ*PSC*)^28^, the (¬*GBM_ce_*) genes, and the (¬*GBM_cf_*) genes ((Equation (3))). A BF ≥6 corresponds to a posterior probability of ∼90% that a gene is essential in a given cell line^26^. Genes that met this strict BF threshold in at least 85%^26^ of GBM cell lines (that passed biological and technical QC) were considered to be ‘GBM core essential’ genes (*GBM_e_*). We defined ‘GBM-specific core essential’ genes as the set difference between the generally-essential genes and the ‘GBM core essential’genes (Equation (4)). As expected, there were no ‘GBM-specific core essential’ genes. We next identified ‘GBM core fitness’ (*GBM_cf_*) and ‘GBM commonly essential’ (*GBM_ce_*) genes in our curated GBM cell line dataset using the ADaM and FiPer algorithms respectively, both of which use a less strict threshold for essentiality. We defined ‘GBM-specific commonly essential’ genes as the set difference between the generally-essential genes and the union of these gene sets (Equation (5)). The intersection of these gene lists is visualised in Fig 1A and B.

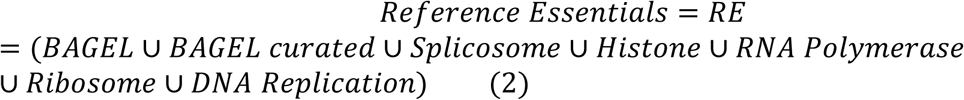

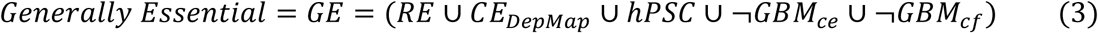

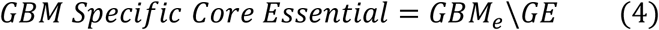

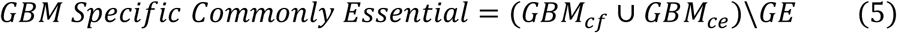

### Annotating cell lines with candidate genetic and transcriptomic biomarkers that are relevant to GBM

#### Annotating cell lines based on the presence of driver mutations and copy number aberrations that are commonly seen in GBM

We identified the following genetic driver mutations in GBM^11–13,91^:

- Loss-of-function mutations, or copy-number deletion of: *TP53, PTEN, NF1, RB1, CDKN2A, CDKN2B, SPTA1, PIK3R1,* and *QKI*.
- Copy-number amplification of: *EGFR, CDK4, PDGFRA, MDM2, MDM4, AKT3, MYCN, SOX2, CDK6, MET, MYC, CCND2, AKT1, ERBB2,* and *PIK3CA*.
- Oncogenic gain-of-function mutations of *EGFR* (e.g. *EGFR^vIII^*) or *PIK3CA*.

We annotated each GBM-cell line screened with these features based on whole-exome sequencing (WXS) or whole-genome sequencing (WGS) data (depending on the available source data for each cell line). Data available from each study was heterogeneous (Table 1). In all cases, the original author-processed variant calls were used. Genetic events with similar anticipated consequences at the proteomic level were grouped into binary features. For example, both copy number deletion and loss of function mutations in PTEN were categorised as ‘PTEN-loss.’ I make the following assumption in identifying such driver events: Finding a loss-of-function genetic event (SNP or copy-number deletion) in a gene known to be a canonical GBM driver in a cell line increases the likelihood that it is a true driver in that GBM cell line, rather than an insignificant passenger mutation. a deleterious genetic event (SNP or copy-number deletion) is found in a gene known canonically to drive GBM, it is more likely this event is a bona fide loss-of-function driver event in that GBM cell line than that it is an irrelevant passenger mutation.

This assumption can be made because I am only looking for established drivers in GBM, not trying to identify novel ones.

We confirmed that each putative oncogenic gain of function mutation was unlikely to be a non-oncogenic passenger mutation using the Ensemble Variant Effect Predictor^92^.

### Annotating cell lines based on enrichment for gene expression signatures that have been described in GBM

Analysis of bulk GBM tissue-derived RNA-seq data identified four GBM subtypes: Mesenchymal, Proneural, Neural and Classical^13^. After subsequent work, the Neural subtype was omitted from these subtypes^14^. Multiple studies have subsequently used single-cell RNA-seq data to demonstrate cell state heterogeneity within individual GBM tumours^15,17,93,94^. Three cell states were found to be anchored to neurodevelopmental cell types (neural progenitor-like; NPC-like, oligodendrocyte progenitor-like; OPC-like, and astrocyte-like; AC-like), and a fourth mesenchymal state (MES-like) that is not anchored in neurodevelopment^15^. Furthermore, these states are thought to be dynamically interchangeable.^15^ An alternative classification describes ‘proliferative/progenitor’ (PPM), ‘neuronal’ (NEU), ‘mitochondrial’ (MTC) and ‘glycolytic/plurimetabolic’ (GPM) cell-states^17^. Another classification describes GBM cells existing on an axis of differentiation between Glioma stem-cell like (G-STEM) to differentiated GBM cells (DGCs)^16^. Finally, Glioma stem cells have been described to exist on a gradient between a ‘developmental’ and an ‘injury response’ state^5^. We used a modified version of the single sample gene set enrichment (ssGSEA) algorithm^14^, extended to analyse the bulk RNA-seq data for all cell lines for enrichment in the gene sets described above.

### Biological quality control - removal of models that were not sufficiently ‘Glioblastoma’- like

Traditional immortalised cell lines of GBM have been criticised for not accurately recapitulating human disease cell states, in part because they are affected by iterative genetic/ transcriptomic drift, particularly towards the Mesenchymal state^32^. We considered cell lines that did were not significantly enriched (<0.05) for any of the above transcriptomic signatures, to be insufficiently representative of human GBM, and omitted them from our analyses. Additionally, we excluded any cell lines that were IDH-mutant, because modern definitions of primary GBM specify it must be IDH-wt^95^.

### Batch correction of RNA-seq counts with CombatSeq

The raw RNA-seq data was not available for every cell line that we included. Where raw RNA-seq count data was available, we ran ComBat-seq^96^ with default settings. ComBat-seq uses a negative binomial regression model to remove batch effects and recover the biological signals within, while retaining the integer nature of RNA count data^96^. We performed “Trimmed mean of M-values” (TMM) normalisation^97^ to facilitate within and between sample comparison. We used this normalised data to create plots which compare RNA expression against gene essentiality (Fig 8).

### Annotation of cell lines based on MGMT expression

MGMT promotor methylation status is predictive of overall survival in GBM treated with temozolomide^18,30^. It was therefore important to consider this as a feature in our modelling. Because methylation data was not available for all cell lines, we instead used MGMT RNA expression in place of MGMT-promotor methylation status. Biologically, we would expect that methylation of the MGMT-promotor will lead to lower expression of MGMT RNA, and subsequently reduced translation into functional MGMT protein. In human GBM samples, MGMT promotor methylation correlates with low MGMT RNA expression^98^. Because raw RNA-seq count data was not available for all cell lines (in some cell lines we only had access to the normalised expression results), it was not possible to make a batch-corrected RNA-expression dataset for all cell lines included in the modelling. We, therefore, took the following approach to dichotomise MGMT expression as ‘expressed’ or ‘suppressed’. For each cell line, RNA expression follows a bimodal distribution. The first, larger, peak is associated with genes of no (or negligible) expression, and the second smaller peak, with expressed genes. We considered MGMT expression to be suppressed if it was below the sum of the lower mode plus 75% of the difference between the first and second modes.

### Annotation of cell lines based on TERT expression

Telomere lengthening is required for cancer cell survival, which can be achieved in various ways, one of which is via telomerase activity. Mutations in the Telomerase Reverse Transcriptase promoter (TERTp) occur in 83% of GBM^19^. These mutations increase recruitment of GABP, a transcription factor, which results in increased TERT expression and subsequent Telomerase activity^99^. Whole exome sequencing (WXS) is a cost-effective way to sequence only the protein-coding regions of a genome, which omits additional components such as promotor regions^100^. Conversely, whole genome sequencing (WGS) sequences the whole genome, which does include promotor regions. Some cell lines included in this study only had WXS data available; WGS data was not available. It was therefore not possible to comprehensively annotate every cell line based on TERTp mutation. Despite this limitation, we felt that TERT status was important to include as a candidate biomarker in our analysis, because of how commonly it occurs in GBM. TERT expression is repressed in most human somatic cells, silencing telomerase activity^101^. We reasoned, therefore, that annotating cell lines based on TERT RNA expression (dichotomised as either ‘expressed’ or ‘suppressed’) may be a sufficient alternative to annotating TERTp mutation. To do this, we used the same method as described to dichotomise MGMT expression.

### Annotation of cell lines based on the expression of GBM relevant paralogous genes

We reviewed the literature to identify which genes are known to be hypermethylated in GBM^20,21^. We used the PANTHER database to identify which of these had known paralogues to genes that were included within our Targeted Learning analysis. Additionally, we included genes that are found within the various transcriptomic signatures described above. We did so because genes that are upregulated in one cell-state / subtype, are by definition downregulated in at least one of the counterpart cell-states / subtypes within that classification system.

### Application of Targeted Learning (Super Learner and Targeted Maximum Likelihood Estimation) to identify context-dependent gene essentiality

#### Identifying genes ‘sometimes essential’ in GBM genes (SEGs) in GBM while avoiding selective inference

To avoid selective inference^40^, we split our dataset and restricted our Targeted Learning analysis to the above genes which had high essentiality (BF of ≥ 6) in at least two of the 14 cell lines that passed biological/ technical QC, but that were not eligible for modelling (either due to missing genetic annotation or cell line duplication) Fig 3a. We then excluded genes within the Generally-Essential group. Finally we arrived at 536 candidate SEGs for which to perform Targeted Learning Fig 3b

#### Defining a SL

Super Learning is comprehensively described elsewhere^38^. Briefly, a Super Learner (SL) is a ‘stack’ of different machine learning algorithms tasked with the same estimation problem. To guard against overfitting, we used 20-fold stratified cross-validation (CV) where possible, and leave-one-out (LOO) CV for rarer biomarkers, to train each algorithm in the library^41^. After performing this task, the learners are ranked based on the accuracy of their predictions (here, as measured by the mean-squared error), and an ensemble machine learner is created using a combination of the most accurate algorithms, weighted by their relative success. We then used a non-negative least squares (NNLS) regression meta-learner to generate a continuous ensemble SL^41^.

Our SL library contains the following algorithms which run after an interactions learner and either a random-forest or lasso feature screener: Least absolute shrinkage and selection operator (Lasso) regression (α = 1)^102^, Ridge regression (α = 0)^102^, Elastic-Net regression (α = 0.5)^102^, Highly Adaptive Lasso (HAL)^103^, Random Forest regression^104^, and eXtreme Gradient Boosting (XGBboost)^105^ algorithms Fig 3d. Other than the α values specified for the above algorithms, we did not impose additional hyperparameters for these algorithms. This is the preferred approach during Super Learning, which allows each algorithm’s internal cross-validation to chose optimal hyper parameters.

Certain genomic drivers and transcriptomic states commonly co-occur. For example, EGFR-amplification is often found in cells adopting an AC-like cell state^15^, and is associated with Verhaak Classical GBM, but all three entities are distinct^13^. To encompass this existing knowledge, each algorithm was run with the specification of a curated set of known interactions between biomarkers/ features Fig 3c).

#### TMLE

We applied this SL in a TMLE framework to double-robustly estimate the ‘average treatment effect’ *ATE* as follows, where *A* = 1 indicates that the biomarker of interest is present in the cell line, and *A* = 0 indicates that it is absent on outcome *Y* (gene essentiality measured as a normalised BF) accounting for covariates *W* (all other candidate biomarkers, sgRNA library, cell line type, and study of origin):

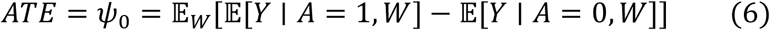

Targeted Learning performs two fits via the SL. Firstly the outcome (*Y*) is fitted as a function of the biomarker (*A*) and covariates (*W*). Secondly, a propensity score of *A* is fitted: *p(A*|*W*). Double robustness of *ATE* means that as long as at least one of these estimates is consistent, then the *ATE* will be estimated consistently with coverage of ground-truth. We also similarly estimated the treatment-specific mean (*TSM*):

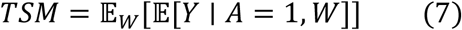

With this we generated the *ATE* and *TSM* for each curated biomarker in each gene, allowing us to compare between genes Fig 3F. Asymptotically valid confidence intervals based on the TMLE were used to derive p-values, and Benjamini-Hochberg corrected p-values based on the number of biomarkers examined per gene. We used the tmle3 package^106^ to implement this.

#### Validation of Targeted Learning finite sample statistical robustness using asymptotic statistical analyses

We validated the robustness of our asymptotic statistical analysis (SL+TMLE), ensuring that our 95% confidence intervals captured the *ATE* derived using finite-sample statistical analysis.Fig 10 The finite-sample analysis technique we used was the HAL-TMLE-Bootstrap^107^. In this analysis, for each candidate biomarker, we performed 5000 bootstrapped TMLE calculations using the following SL The SL contained a random forest variable screener, followed by the HAL. To improve computational run-time, we ran HAL with the following recommended settings^108^: the smoothness of the basis functions was set to 0, The highest order of interaction terms was set to 3, and the maximum number of knot points was set to (50, 25, 10). We performed this analysis on the three genes of interest (VRK1, RPP25L and WWTR1; Fig 10A, B, and C respectively) in which there were known, orthogonally validated target-biomarker pairs (VRK2-expression^109,109^, RPP25-expression^45^ and EGFR^mut/amp44^ respectively). We demonstrate the changing mean *ATE* with *m* increasing bootstraps, which stabilises above 1000 bootstraps Fig 10A, B, and C left panels). The standard deviation also similarly stabilises as *m* increases (data not shown). We took the mean of all 5000 bootstraps to calculate the finite-sample derived *ATE*. We ranked the bootstrapped *ATE* values and extracted the 2.5% and 97.5% values to infer 95% confidence intervals.

**Fig 10:**
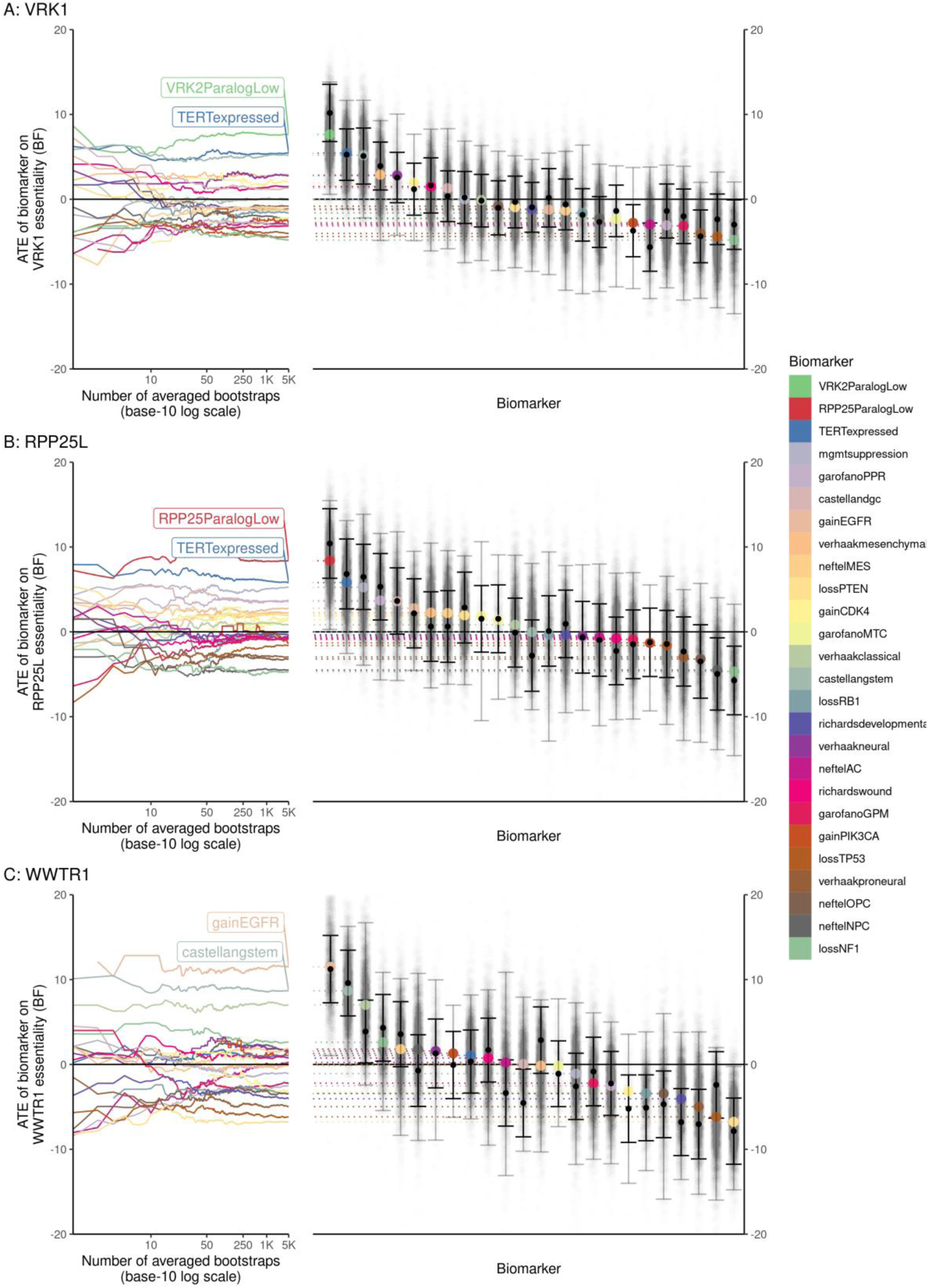
Validation of asymptotic statistical analysis using finite-sample analysis. We chose three targets - VRK1 (A), RPP25L (B) and WWTR1 (C) - which already had orthogonally validated biomarkers (VRK2-expression^42,112^, RPP25-expression^45^ and EGFR^mut/amp^^44^ respectively) with which to check the robustness of our statistical analyses. We wanted to confirm that the *ATE* derived by (computationally intensive, but theoretically more accurate) finite sample statistical analysis (bootstrapped-HAL-TMLE^107^ (see Methods)) was within the confidence intervals of our main (asymptotic) statistical analysis (SL-TMLE). In each row, the left graph shows the averaged *ATE* derived from increasing numbers of bootstraps, on a Log-10 scale. In all cases, the finite sample derived *ATE* was within the confidence intervals of our asymptotic analysis, demonstrating the robustness of our approach. Above 1000 bootstraps these values become stable. In each row, the right graph shows the SL-TMLE derived *ATE* ± confidence intervals (black), the corresponding bootstrap-(HAL-TMLE) derived *ATE* (coloured by biomarker), the 95% confidence intervals of bootstrap-(HAL-TMLE) (translucent grey line), and the individual values of each bootstrap-HAL-TMLE derived *ATE* (translucent grey dots). The y-axis for the left and right graphs on each row are identical, allowing horizontal, appropriately coloured, dotted lines to match up the final bootstrap-(HAL-TMLE) derived *ATE* (left graphs) with its relation to the SL-TMLE derived *ATE* (right graphs)

## Acknowledgements

MTF was supported by the ECAT CRUK PhD Fellowship scheme, and a CRUK Brain Tumour Centre of Excellence Award (C157/A27589). CAS was supported by MRC core funding to the MRC Human Genetics Unit (MRC grant MC_UU_00007/16). AK was supported by the XDF programme from the University of Edinburgh and the Medical Research Council (MC_UU_00009/2). NOC and MF were supported by a CRUK/Brain Tumour Charity funded Brain Tumour Award (C42454/A28596). SVB is supported by a University of Edinburgh Chancellor’s Fellowship from the University of Edinburgh. AE and AK are each supported by a Langmuir Talent Development Fellowship from the Institute of Genetics and Cancer, and a philanthropic donation from Hugh and Josseline Langmuir. We would like to thank Charlie Gourley (ECAT CRUK Supervisor to MTF), members of the Semple lab (Stuart Aitken, Graeme Grimes, Alison Meynert, Thomas Parry, Murray Wham), Lana Talmane, Becka Hughes, Alison Munro, Steve Pollard, and Al Hamden. Thank you to Jan Irvine, Laura Wood, and Jo Ness for administrative support. Thank you to Graham Macleod, Stephane Angers, and Peter Dirks for sharing raw data.

## Author contributions

- AE: Application of statistical, mathematical, computational, or other formal techniques to analyze or synthesize study data. Development or design of methodology; creation of models.
- AK: Introduced, conceptualised and supervised application of Targeted Learning methodology to this work. Application of statistical, mathematical, computational, or other formal techniques to analyze or synthesize study data. Development or design of methodology; creation of models. Preparation, creation and/or presentation of the published work by those from the original research group, specifically critical review, commentary or revision – including pre-or postpublication stages.
- CS: Oversight and leadership responsibility for the research activity planning and execution, including mentorship external to the core team. Preparation, creation and/or presentation of the published work by those from the original research group, specifically critical review, commentary or revision – including pre-or postpublication stages.
- MF: Oversight and leadership responsibility for the research activity planning and execution, including mentorship external to the core team. Acquisition of the financial support for the project leading to this publication. Preparation, creation and/or presentation of the published work by those from the original research group, specifically critical review, commentary or revision – including pre-or postpublication stages.
- MTF: Preparation, creation and/or presentation of the published work, specifically writing the initial draft. Application of statistical, mathematical, computational, or other formal techniques to analyze or synthesize study data. Acquisition of the financial support for the project leading to this publication. Programming, software development; designing computer programs; implementation of the computer code and supporting algorithms; testing of existing code components. Development or design of methodology; creation of models. Preparation, creation and/or presentation of the published work, specifically visualization/ data presentation. Management activities to annotate (produce metadata), scrub data and maintain research data (including software code, where it is necessary for interpreting the data itself) for initial use and later reuse.
- NC: Oversight and leadership responsibility for the research activity planning and execution, including mentorship external to the core team. Acquisition of the financial support for the project leading to this publication. Preparation, creation and/or presentation of the published work by those from the original research group, specifically critical review, commentary or revision – including pre-or postpublication stages.
- PMB: Oversight and leadership responsibility for the research activity planning and execution, including mentorship external to the core team. Preparation, creation and/or presentation of the published work by those from the original research group, specifically critical review, commentary or revision – including pre-or postpublication stages.
- SVB: Introduced, conceptualised and supervised application of Targeted Learning methodology to this work. Application of statistical, mathematical, computational, or other formal techniques to analyze or synthesize study data. Development or design of methodology; creation of models. Preparation, creation and/or presentation of the published work by those from the original research group, specifically critical review, commentary or revision – including pre-or postpublication stages.

## Ethics declarations

MTF: None.

AE: None.

MF: None.

PMB: None.

AK: None.

SVB: None.

NC: is a scientific advisory board member and shareholder in Amplia Therapeutics Ltd (Melbourne, Australia); is and founder, shareholder, and management consultant of PhenoTherapeutics Ltd (Edinburgh, UK); is Director of Ther-IP Ltd (Edinburgh); and is Director of Edinburgh Innovations Ltd.

CS: None.

## Supplementary Figures

**Supplementary Fig 1.**
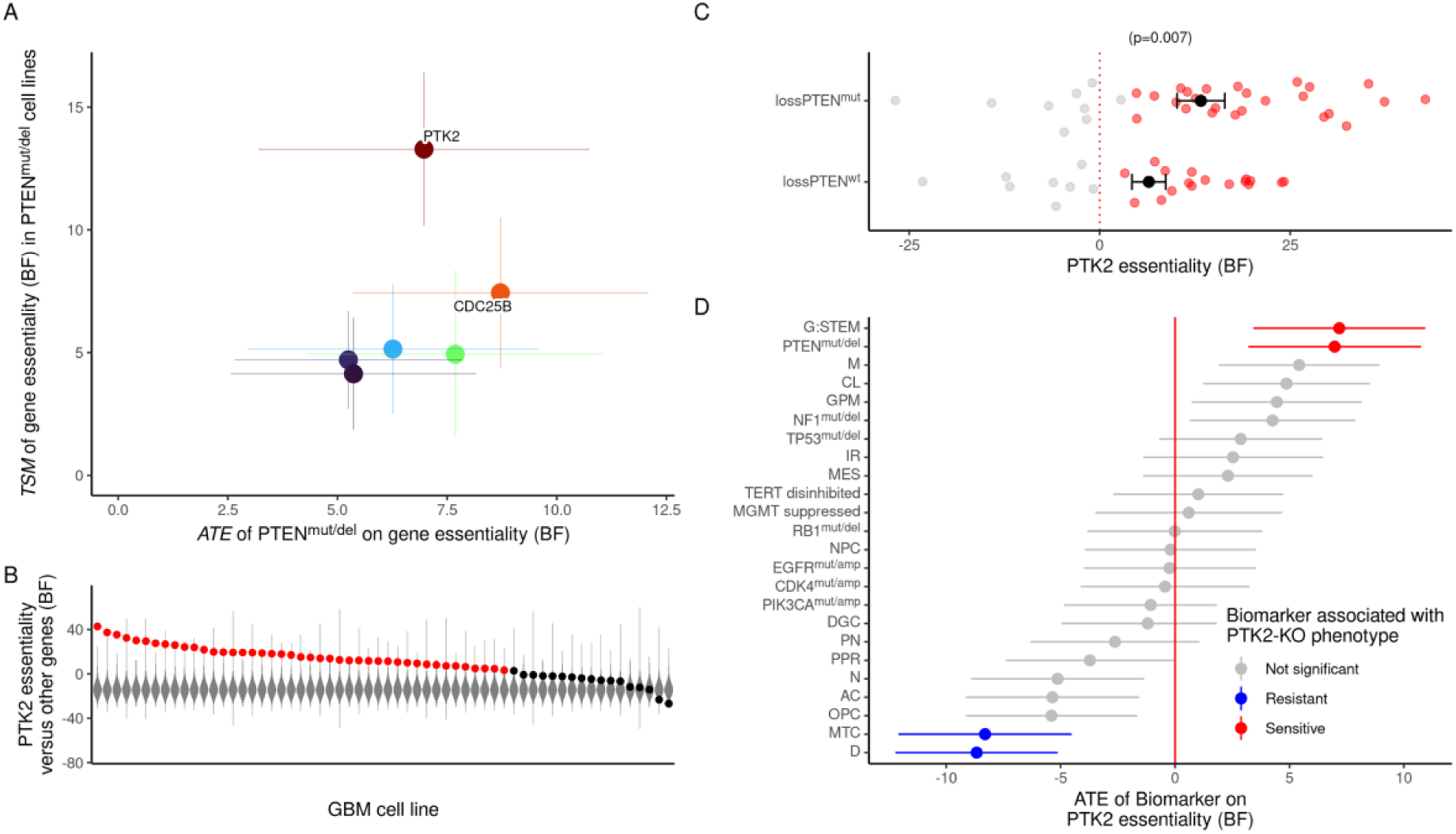
Extended Data Fig 1 PTEN^mut^ is a potential biomarker for PTK2 (FAK) essentiality. The PTK2 gene (also known as FAK), which encodes Focal Adhesion kinase (FAK), was essential in 85% of GBM cell lines. PTEN^mut^ GBM cell lines were more vulnerable to FAK-knockout than PTEN-wildtype (PTEN^wt)^ cell lines This is mechanistically feasible because functionally normal PTEN protein is reported to dephosphorylate FAK^113^. If true, it follows that loss of functioning PTEN protein (as seen commonly in GBM either via loss-of-function mutation or copy-number deletion events) may disinhibit, and in time may endow a dependency on FAK. Further research is needed to experimentally validate this in GBM, but we note that PTEN^mut^ Uterine cancer cell lines were more sensitive to pharmacological FAK-inhibition than corresponding PTEN^wt^ cell lines^114^. This is potentially important, because PTEN^mut^ is found in 23% of GBM samples^12^, and because FAK inhibitors (while well-tolerated in Phase II human trials^115^) require biomarkers of therapeutic efficacy^116^ Abbreviations as per Fig 6.

**Supplementary Fig 2.**
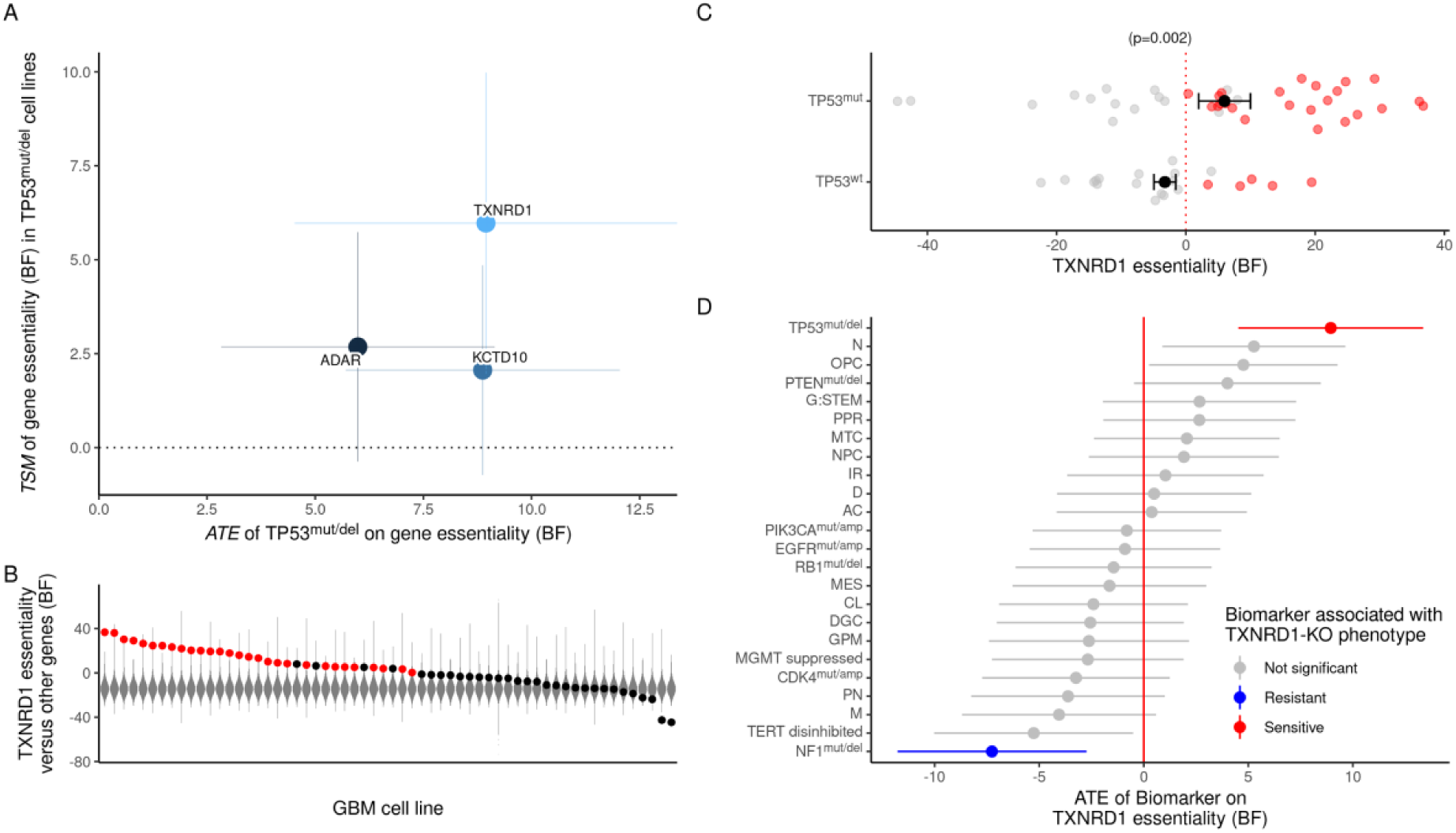
Extended Data Fig 2 TP53^mut^ is a potential biomarker for TXNRD1 essentiality. TXNRD1, which encodes Thioredoxin reductase-1, was an essential gene in 63% of GBM cell lines. GBM cell lines with a loss of function mutation in TP53 (TP53^mut^) were more vulnerable to TXNRD1-knockout than TP53-wildtype (TP53^wt^) GBM cell lines. This is potentially important because TP53^mut^ is seen in 25% of GBM tumours^12^, and because Thioredoxin reductase-1 inhibitors (including the approved, repurposable drug Auranofin) are candidates for anticancer therapy^117^. Although TP53^mut^ has not been described as a biomarker of TXNRD1 essentiality in GBM specifically, we note that it has been described as a biomarker for Auranofin treatment in both non-small-cell lung-cancer^118^ and B-cell lymphoma^119^. Abbreviations as per Fig 6.

**Supplementary Fig 3.**
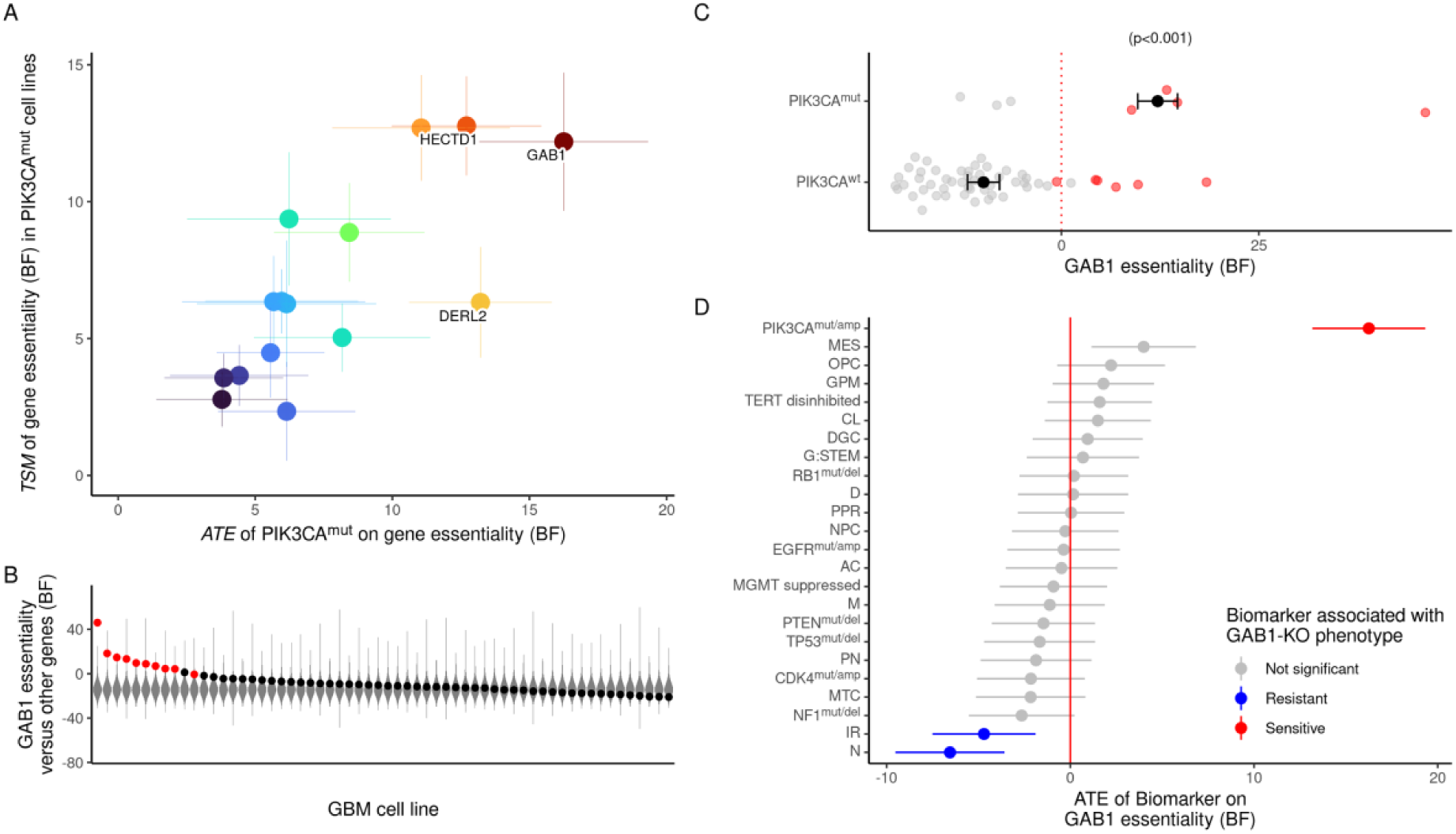
Extended Data Fig 3 PIK3CA^mut^ is a potential biomarker for GAB1 essentiality. [a] We looked at the genes in which PIK3CA mutation was found to have a significant positive impact on *ATE*. GAB1 was the gene associated with the highest *ATE* of PIK3CA mutation [b] GAB1, which encodes GRB2-associated-binding protein 1, was an essential gene in 23% of GBM cell lines. [c] Using a simple univariate analysis GAB1 essentiality scores were higher p<0.001, and GAB1 was more commonly classified as an essential gene (red) than as a non-essential gene (grey) p=0.065. [d] We estimated the *ATE* of multiple biomarkers on GAB1 essentiality; PIK3CA mutation was most associated with increased GAB1 essentiality. NF1 mutation was associated with GAB1 knockout resistance. This putative biomarker pair is potentially useful because PIK3CA is mutated in 10% of GBM^11^, and the GAB1 gene product is amenable to small molecule inhibition^120^. The relationship is mechanistically plausible because GAB1 enables the formation of active PIK3, through the recruitment of the regulatory subunit (p85, encoded by PIK3R1) thus activating the catalytic subunit (p110, encoded by PIK3CA)^121^. Thus, suppression of GAB1 may reduce the activity of oncogenically-mutated PIK3CA on PIK3 activity. However, Further experimental studies are required. Abbreviations as per Fig 6.

## Supplementary Tables

**Supplementary Table S1.** CRISPR datasets from multiple studies were interrogated which included 83 GBM cell lines that were considered for inclusion. From these, duplicates and low quality cell lines, and low-quality CRISPR screens were discarded. This culminated in 74 models available for pan-Glioblastoma / context-agnostic analysis, 60 for context-dependent modelling, and 61 for synergist or paralog analyses.

| Author | Year | Library name<br>(addgene) | Cas9<br>vector<br>addgene | Cell model<br>type | Endpoint<br>time | CRISPR screen<br>data source | Cell line<br>annotation source | Number of GBM<br>models available<br>in dataset, [passed<br>QC, used in TL,<br>Raw RNA-seq<br>data available] |
| --- | --- | --- | --- | --- | --- | --- | --- | --- |
| Toledo et al<sup>7</sup> | 2015 | GeCKO v1<sup>75,122</sup> | #52961 | GSC | 21 days | GSE70038<sup>123</sup> | Toledo et al<sup>7</sup><br>supplementary<br>material | 2 [1, 1, 0] |
| Behan et al<sup>4</sup> | 2019 | Yusa (#67989)<sup>72,73</sup> | #68343 | Immortalised | 14 days | Project Score<sup>124</sup> | DepMap 20q3<sup>69</sup> | 27 [23, 18, 23] |
| Macleod et al<sup>6</sup> | 2019 | TKOv1<br>(#1000000069)<sup>66,76/</sup><br>TKOv3 (#125517)<sup>26,77</sup> | #73310 | GSC | 10-14<br>doublings | Author<br>correspondance | Author<br>correspondance | 10 [9, 5, 7] |
| Dempster et al<sup>125</sup> & Meyers et al<sup>8</sup> | 2020 | AVANA<sup>68,126</sup> | #96924 | Immortalised | 21 days | DepMap 20q3<sup>69</sup> | DepMap 20q3<sup>69</sup> | 33 [31, 28, 31] |
| Richards et al<sup>5</sup> | 2021 | TKOv3 (#125517)<sup>26,77</sup> | #73310 | GSC | 10-14<br>doublings | Author<br>correspondance | Author<br>correspondance | 11 [10, 8, 0] |

**Supplementary Table S2.**
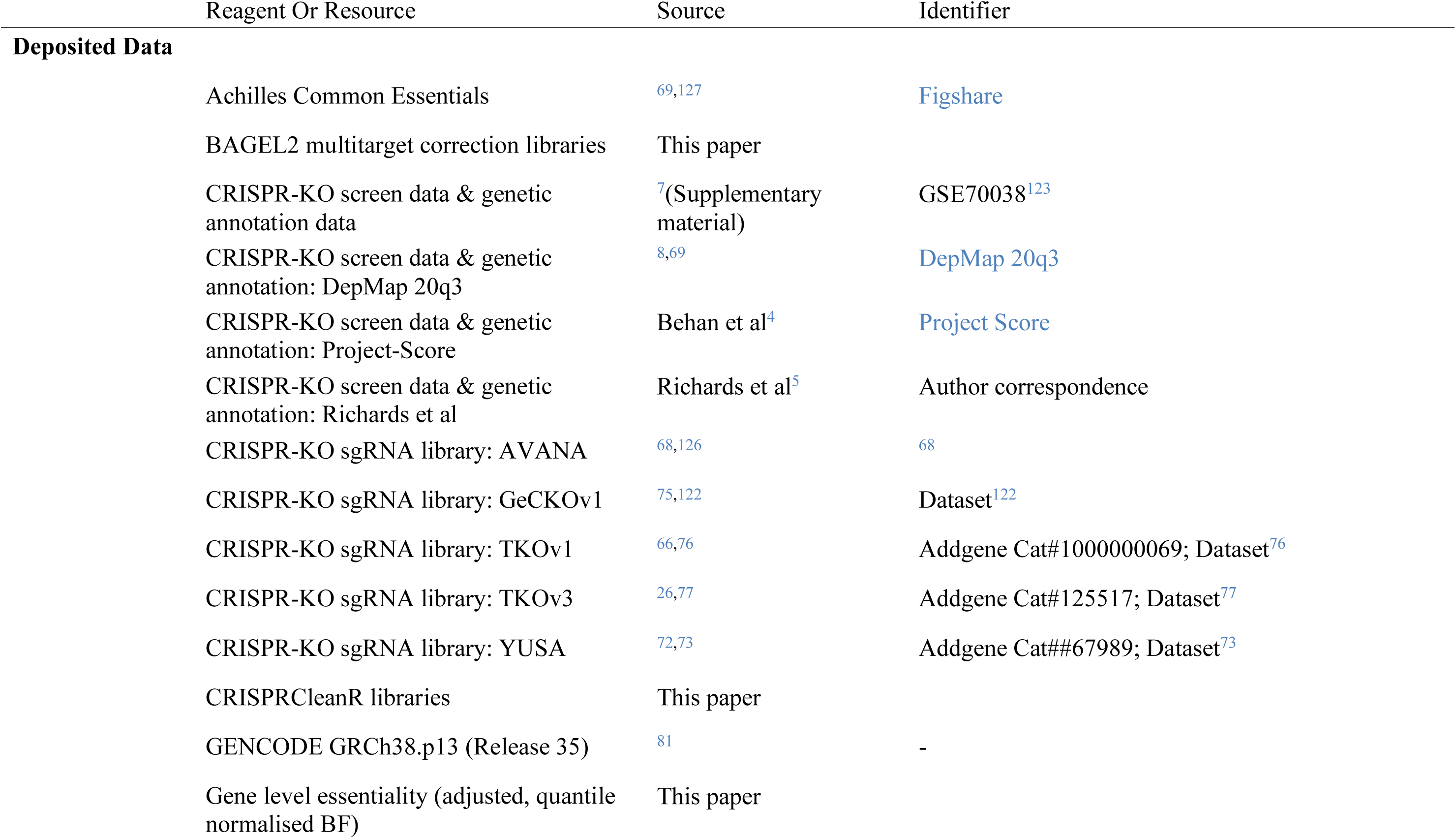
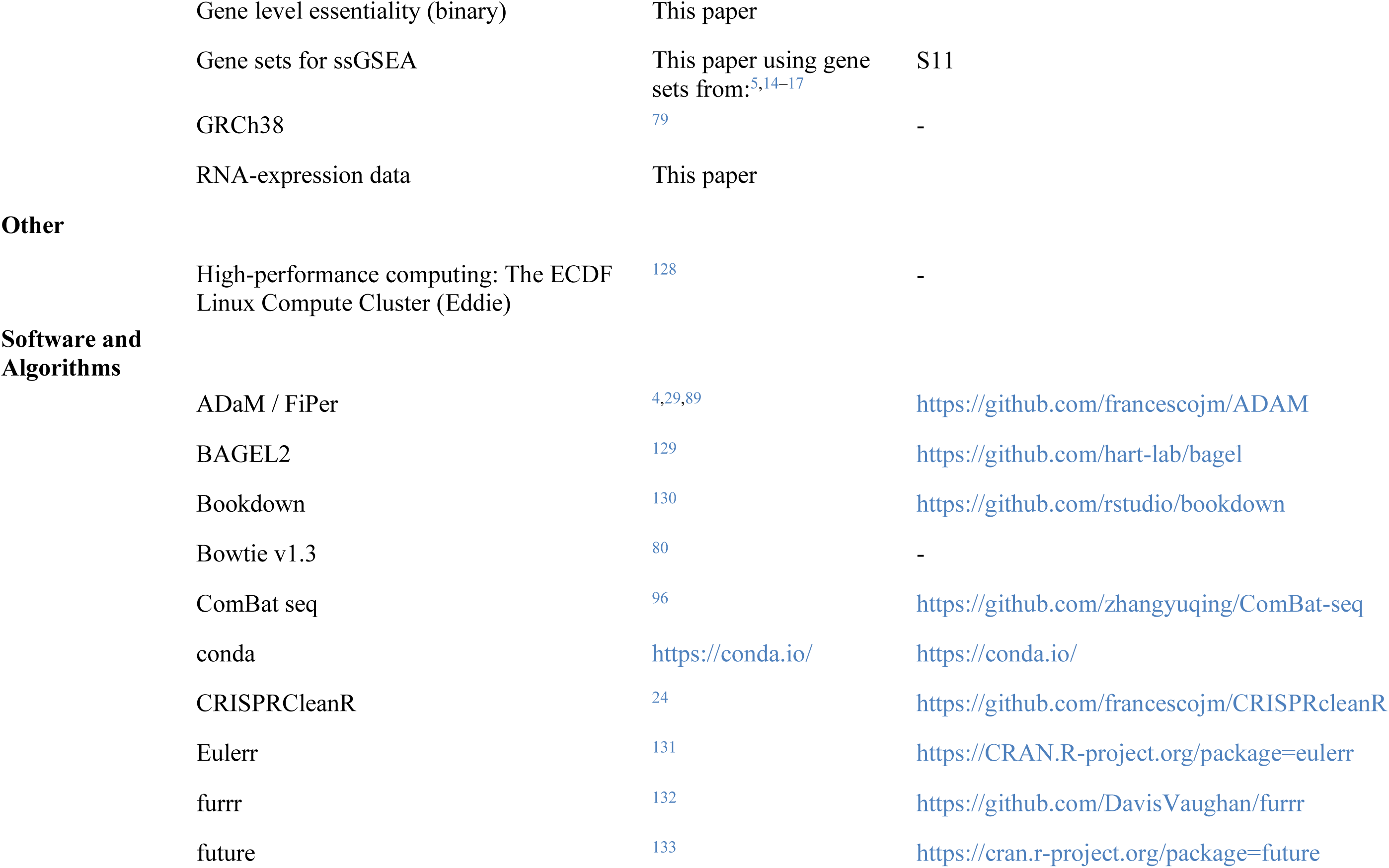
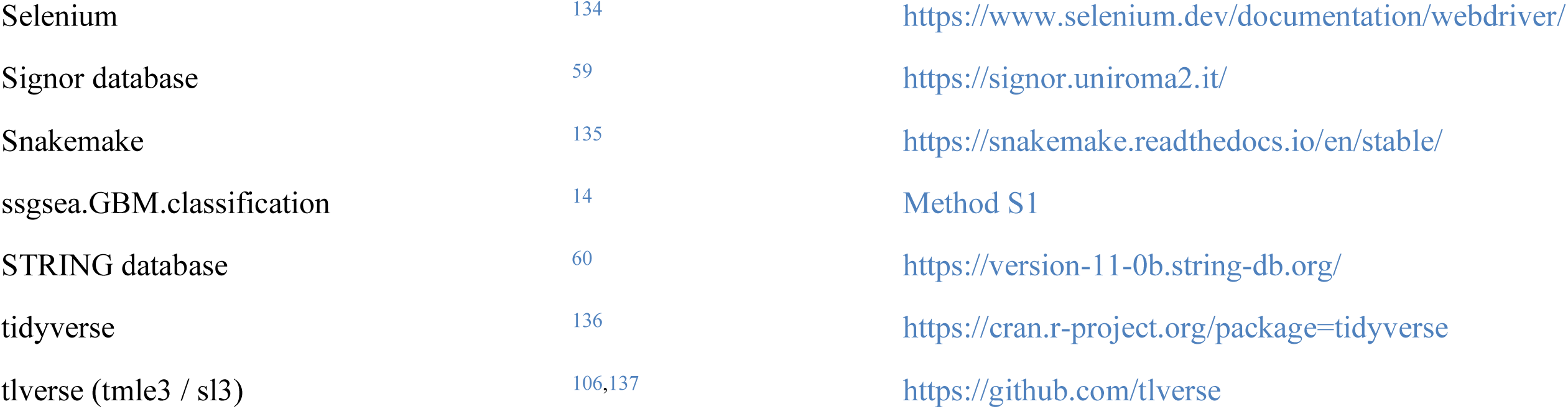
Methods table.

| Reagent Or Resource | Source | Identifier |
| --- | --- | --- |
| **Deposited Data** |  |  |
| Achilles Common Essentials | 69,127 | Figshare |
| BAGEL2 multitarget correction libraries | This paper |  |
| CRISPR-KO screen data & genetic annotation data | 7(Supplementary material) | GSE70038<sup>123</sup> |
| CRISPR-KO screen data & genetic annotation: DepMap 20q3 | 8,69 | DepMap 20q3 |
| CRISPR-KO screen data & genetic annotation: Project-Score | Behan et al<sup>4</sup> | Project Score |
| CRISPR-KO screen data & genetic annotation: Richards et al | Richards et al<sup>5</sup> | Author correspondence |
| CRISPR-KO sgRNA library: AVANA | 68,126 | 68 |
| CRISPR-KO sgRNA library: GeCKOv1 | 75,122 | Dataset<sup>122</sup> |
| CRISPR-KO sgRNA library: TKOv1 | 66,76 | Addgene Cat#1000000069; Dataset<sup>76</sup> |
| CRISPR-KO sgRNA library: TKOv3 | 26,77 | Addgene Cat#125517; Dataset<sup>77</sup> |
| CRISPR-KO sgRNA library: YUSA | 72,73 | Addgene Cat##67989; Dataset<sup>73</sup> |
| CRISPRCleanR libraries | This paper |  |
| GENCODE GRCh38.p13 (Release 35) | 81 | - |
| Gene level essentiality (adjusted, quantile normalised BF) | This paper |  |

|  |  |  |  |
| --- | --- | --- | --- |
|  | Gene level essentiality (binary) | This paper |  |
|  | Gene sets for ssGSEA | This paper using gene sets from:<sup>5,14–17</sup> | S11 |
|  | GRCh38 | <sup>79</sup> | - |
|  | RNA-expression data | This paper |  |
| **Other** |  |  |  |
|  | High-performance computing: The ECDF Linux Compute Cluster (Eddie) | <sup>128</sup> | - |
| **Software and Algorithms** |  |  |  |
|  | ADaM / FiPer | <sup>4,29,89</sup> | https://github.com/francescojm/ADAM |
|  | BAGEL2 | <sup>129</sup> | https://github.com/hart-lab/bagel |
|  | Bookdown | <sup>130</sup> | https://github.com/rstudio/bookdown |
|  | Bowtie v1.3 | <sup>80</sup> | - |
|  | ComBat seq | <sup>96</sup> | https://github.com/zhangyuqing/ComBat-seq |
|  | conda | https://conda.io/ | https://conda.io/ |
|  | CRISPRCleanR | <sup>24</sup> | https://github.com/francescojm/CRISPRcleanR |
|  | Eulerr | <sup>131</sup> | https://CRAN.R-project.org/package=eulerr |
|  | furrr | <sup>132</sup> | https://github.com/DavisVaughan/furrr |
|  | future | <sup>133</sup> | https://cran.r-project.org/package=future |

Supplementary Table S2
|  |  |  |
| --- | --- | --- |
| Selenium | 134 | https://www.selenium.dev/documentation/webdriver/ |
| Signor database | 59 | https://signor.uniroma2.it/ |
| Snakemake | 135 | https://snakemake.readthedocs.io/en/stable/ |
| ssgsea.GBM.classification | 14 | Method S1 |
| STRING database | 60 | https://version-11-0b.string-db.org/ |
| tidyverse | 136 | https://cran.r-project.org/package=tidyverse |
| tlverse (tmle3 / sl3) | 106,137 | https://github.com/tlverse |

## Notes

### Competing Interest Statement

MTF, AE, MF, PMB, AK, SVB, CS: None.
NC: is a scientific advisory board member and shareholder in Amplia Therapeutics Ltd (Melbourne, Australia); is and founder, shareholder, and management consultant of PhenoTherapeutics Ltd (Edinburgh, UK); is Director of Ther-IP Ltd (Edinburgh); and is Director of Edinburgh Innovations Ltd.

## References

1. Stupp, R. et al. Radiotherapy plus concomitant and adjuvant temozolomide for glioblastoma. The New England Journal of Medicine 352, 987–996 (2005).

2. Lord, C. J. & Ashworth, A. PARP inhibitors: Synthetic lethality in the clinic. Science 355, 1152–1158 (2017).

3. Huang, A., Garraway, L. A., Ashworth, A. & Weber, B. Synthetic lethality as an engine for cancer drug target discovery. Nature Reviews Drug Discovery 1–16 (2019) doi:10.1038/s41573-019-0046-z.

4. Behan, F. M. et al. Prioritization of cancer therapeutic targets using CRISPR-Cas9 screens. Nature (2019) doi:10.1038/s41586-019-1103-9.

5. Richards, L. M. et al. Gradient of Developmental and Injury Response transcriptional states defines functional vulnerabilities underpinning glioblastoma heterogeneity. Nature Cancer 1–17 (2021) doi:10.1038/s43018-020-00154-9.

6. MacLeod, G. et al. Genome-Wide CRISPR-Cas9 Screens Expose Genetic Vulnerabilities and Mechanisms of Temozolomide Sensitivity in Glioblastoma Stem Cells. Cell Reports 27, 971–986.e9 (2019).

7. Toledo, C. M. et al. Genome-wide CRISPR-Cas9 Screens Reveal Loss of Redundancy between PKMYT1 and WEE1 in Glioblastoma Stem-like Cells. Cell Reports 13, 2425–2439 (2015).

8. Meyers, R. M. et al. Computational correction of copy number effect improves specificity of CRISPR-Cas9 essentiality screens in cancer cells. Nature Genetics 49, 1779– 1784 (2017).

9. Louis, D. N., et al. The 2016 World Health Organization Classification of Tumors of the Central Nervous System: A summary. Acta Neuropathologica 131, 803–820 (2016).

10. Gonzalez Castro, L. N. & Wesseling, P. The cIMPACT-NOW updates and their significance to current neuro-oncology practice. Neuro-Oncology Practice 8, 4–10 (2021).

11. Brennan, C. W. et al. The Somatic Genomic Landscape of Glioblastoma. Cell 155, 462–477 (2013).

12. Martínez-Jiménez, F. et al. A compendium of mutational cancer driver genes. Nature Reviews Cancer 20, 555–572 (2020).

13. Verhaak, R. G. W. et al. Integrated genomic analysis identifies clinically relevant subtypes of glioblastoma characterized by abnormalities in PDGFRA, IDH1, EGFR, and NF1. Cancer Cell 17, 98–110 (2010).

14. Wang, Q. et al. Tumor Evolution of Glioma-Intrinsic Gene Expression Subtypes Associates with Immunological Changes in the Microenvironment. Cancer Cell 32, 42–56.e6 (2017).

15. Neftel, C. et al. An Integrative Model of Cellular States, Plasticity, and Genetics for Glioblastoma. Cell 178, 835–849.e21 (2019).

16. Castellan, M. et al. Single-cell analyses reveal YAP/TAZ as regulators of stemness and cell plasticity in Glioblastoma. Nature Cancer 2, 174–188 (2021).

17. Garofano, L. et al. Pathway-based classification of glioblastoma uncovers a mitochondrial subtype with therapeutic vulnerabilities. Nature Cancer 1–16 (2021) doi:10.1038/s43018-020-00159-4.

18. Hegi, M. E. et al. MGMT gene silencing and benefit from temozolomide in glioblastoma. The New England Journal of Medicine 352, 997–1003 (2005).

19. Killela, P. J. et al. TERT promoter mutations occur frequently in gliomas and a subset of tumors derived from cells with low rates of self-renewal. Proceedings of the National Academy of Sciences of the United States of America 110, 6021–6026 (2013).

20. Ferreira, W. A. S. et al. An update on the epigenetics of glioblastomas. Epigenomics 8, 1289–1305 (2016).

21. Wang, W. et al. Bioinformatic analysis of gene expression and methylation regulation in glioblastoma. Journal of Neuro-Oncology 136, 495–503 (2018).

22. Fujisawa, H. et al. Loss of Heterozygosity on Chromosome 10 Is More Extensive in Primary (De Novo) Than in Secondary Glioblastomas. Laboratory Investigation 80, 65–72 (2000).

23. Laan, M. J. laanvan der & Rubin, D. Targeted Maximum Likelihood Learning. The International Journal of Biostatistics 2, (2006).

24. Iorio, F. et al. Unsupervised correction of gene-independent cell responses to CRISPR-Cas9 targeting. BMC genomics 19, 604 (2018).

25. Improved analysis of CRISPR fitness screens and reduced off-target effects with the BAGEL2 gene essentiality classifier | bioRxiv.

26. Hart, T. et al. Evaluation and Design of Genome-Wide CRISPR/SpCas9 Knockout Screens. 7, 2719–2727 (2017).

27. Pacini, C. et al. Integrated cross-study datasets of genetic dependencies in cancer. Nature Communications 12, 1661 (2021).

28. Yilmaz, A., Peretz, M., Aharony, A., Sagi, I. & Benvenisty, N. Defining essential genes for human pluripotent stem cells by CRISPR-Cas9 screening in haploid cells. Nature Cell Biology 20, 610–619 (2018).

29. Vinceti, A. et al. CoRe: A robustly benchmarked R package for identifying core-fitness genes in genome-wide pooled CRISPR-Cas9 screens. BMC genomics 22, 828 (2021).

30. McAleenan, A. et al. Prognostic value of test(s) for O6-methylguanine-DNA methyltransferase (MGMT) promoter methylation for predicting overall survival in people with glioblastoma treated with temozolomide. The Cochrane Database of Systematic Reviews 3, CD013316 (2021).

31. Warren, A. et al. Global computational alignment of tumor and cell line transcriptional profiles. Nature Communications 12, 22 (2021).

32. “Malignant Glioma Cell Line - an overview ScienceDirect Topics.” Available: https://www.sciencedirect.com/topics/medicine-and-dentistry/malignant-glioma-cell-line. [Accessed: Feb. 26, 2020]

33. Tsherniak, A. et al. Defining a Cancer Dependency Map. Cell 170, 564–576.e16 (2017).

34. Ito, T. et al. Paralog knockout profiling identifies DUSP4 and DUSP6 as a digenic dependence in MAPK pathway-driven cancers. Nature Genetics 53, 1664–1672 (2021).

35. Lord, C. J., Quinn, N. & Ryan, C. J. Integrative analysis of large-scale loss-of-function screens identifies robust cancer-associated genetic interactions. eLife 9, e58925 (2020).

36. De Kegel, B., Quinn, N., Thompson, N. A., Adams, D. J. & Ryan, C. J. Comprehensive prediction of robust synthetic lethality between paralog pairs in cancer cell lines. Cell Systems 12, 1144–1159.e6 (2021).

37. Viswanathan, S. R. et al. Genome-scale analysis identifies paralog lethality as a vulnerability of chromosome 1p loss in cancer. Nature Genetics 50, 937–943 (2018).

38. Laan, M. J. laanvan der, Polley, E. C. & Hubbard, A. E. Super Learner. Statistical Applications in Genetics and Molecular Biology 6, (2007).

39. laanvan der Laan, M. J. & Starmans, R. J. C. M. Entering the Era of Data Science: Targeted Learning and the Integration of Statistics and Computational Data Analysis. Advances in Statistics 2014, e502678 (2014).

40. Taylor, J. & Tibshirani, R. J. Statistical learning and selective inference. Proceedings of the National Academy of Sciences 112, 7629–7634 (2015).

41. Phillips, R. V., laanvan der Laan, M. J., Lee, H. & Gruber, S. Practical considerations for specifying a super learner. (2022) doi:10.48550/arXiv.2204.06139.

42. Shields, J. A. et al. VRK1 is a Synthetic Lethal Target in VRK2-deficient Glioblastoma. Cancer Research CAN-21–4443 (2022) doi:10.1158/0008-5472.CAN-21-4443.

43. So, J. et al. VRK1 as a synthetic lethal target in VRK2-methylated cancers of the nervous system. JCI Insight (2022) doi:10.1172/jci.insight.158755.

44. Vigneswaran, K. et al. YAP/TAZ Transcriptional Coactivators Create Therapeutic Vulnerability to Verteporfin in EGFR-mutant Glioblastoma. Clinical Cancer Research: An Official Journal of the American Association for Cancer Research 27, 1553–1569 (2021).

45. Köferle, A. et al. Interrogation of cancer gene dependencies reveals paralog interactions of autosome and sex chromosome-encoded genes. Cell Reports 39, 110636 (2022).

46. Campillo-Marcos, I., García-González, R., Navarro-Carrasco, E. & Lazo, P. A. The human VRK1 chromatin kinase in cancer biology. Cancer Letters 503, 117–128 (2021).

47. Moura, D. S. et al. Oncogenic Sox2 regulates and cooperates with VRK1 in cell cycle progression and differentiation. Scientific Reports 6, 28532 (2016).

48. Bulstrode, H. et al. Elevated FOXG1 and SOX2 in glioblastoma enforces neural stem cell identity through transcriptional control of cell cycle and epigenetic regulators. Genes & Development 31, 757–773 (2017).

49. Vinagre, J. et al. Frequency of TERT promoter mutations in human cancers. Nature Communications 4, 2185 (2013).

50. Choi, Y. H., Lim, J.-K., Jeong, M.-W. & Kim, K.-T. HnRNP A1 phosphorylated by VRK1 stimulates telomerase and its binding to telomeric DNA sequence. Nucleic Acids Research 40, 8499–8518 (2012).

51. Gan, H. K., Kaye, A. H. & Luwor, R. B. The EGFRvIII variant in glioblastoma multiforme. Journal of Clinical Neuroscience: Official Journal of the Neurosurgical Society of Australasia 16, 748–754 (2009).

52. Read, W. A Phase 1 / 2 Study of Visudyne (Liposomal Verteporfin) in Persons With Recurrent High Grade EGFR-Mutated Glioblastoma. (2021).

53. Parrish, P. C. R. et al. Discovery of synthetic lethal and tumor suppressor paralog pairs in the human genome. Cell Reports 36, 109597 (2021).

54. Venkataramani, V. et al. Glioblastoma hijacks neuronal mechanisms for brain invasion. Cell 185, 2899–2917.e31 (2022).

55. Aldaz, P. et al. High SOX9 Maintains Glioma Stem Cell Activity through a Regulatory Loop Involving STAT3 and PML. International Journal of Molecular Sciences 23, 4511 (2022).

56. O’Connor, S. A. et al. Neural G0: A quiescent-like state found in neuroepithelial-derived cells and glioma. Molecular Systems Biology 17, e9522 (2021).

57. Tian, R. et al. Genome-wide CRISPRi/a screens in human neurons link lysosomal failure to ferroptosis. Nature Neuroscience 24, 1020–1034 (2021).

58. CRISPRbrain Simple Screen.” Available: https://crisprbrain.org/simple-screen/?screen=Glutamatergic%20Neuron-Survival-CRISPRi. [Accessed: Sep. 12, 2022]

59. Licata, L., et al. SIGNOR 2.0, the SIGnaling Network Open Resource 2.0: 2019 update. Nucleic Acids Research 48, D504–D510 (2020).

60. Szklarczyk, D., et al. STRING V10: Protein-protein interaction networks, integrated over the tree of life. Nucleic Acids Research 43, D447–452 (2015).

61. Dempster, J. M. et al. Gene expression has more power for predicting in vitro cancer cell vulnerabilities than genomics. bioRxiv 2020.02.21.959627 (2020) doi:10.1101/2020.02.21.959627.

62. Iorio, F., et al. A Landscape of Pharmacogenomic Interactions in Cancer. Cell 166, 740–754 (2016).

63. “ATLANTIS.” Cancer Data Science at Broad Institute, Oct. 2021. Available: https://github.com/cancerdatasci/atlantis. [Accessed: Nov. 18, 2021]

64. Jackson, C. M., Choi, J. & Lim, M. Mechanisms of immunotherapy resistance: Lessons from glioblastoma. Nature Immunology 20, 1100–1109 (2019).

65. Venkatesh, H. S. et al. Electrical and synaptic integration of glioma into neural circuits. Nature 1–7 (2019) doi:10.1038/s41586-019-1563-y.

66. Hart, T. et al. High-Resolution CRISPR Screens Reveal Fitness Genes and Genotype-Specific Cancer Liabilities. Cell 163, 1515–1526 (2015).

67. “DepMap: The Cancer Dependency Map Project at Broad Institute.” Available: https://depmap.org/portal/. [Accessed: Mar. 31, 2020]

68. Meyers, R. M., et al. AVANA Library dataset. Nature Genetics (2017).

69. “Depmap 20q3.” Available: doi:10.6084/m9.figshare.12931238.v1. [Accessed: Dec. 17, 2020]

70. Dwane, L. et al. Project Score database: A resource for investigating cancer cell dependencies and prioritizing therapeutic targets. Nucleic Acids Research (2020) doi:10.1093/nar/gkaa882.

71. “Project Score” Available: https://score.depmap.sanger.ac.uk/ [Accessed: 17/12/2020]

72. Tzelepis, K. et al. A CRISPR Dropout Screen Identifies Genetic Vulnerabilities and Therapeutic Targets in Acute Myeloid Leukemia. Cell Reports 17, 1193–1205 (2016).

73. Tzelepis, K., et al. Yusa Library dataset. (2016).

74. Pollard, S. M. et al. Glioma stem cell lines expanded in adherent culture have tumor-specific phenotypes and are suitable for chemical and genetic screens. Cell Stem Cell 4, 568– 580 (2009).

75. Shalem, O., et al. Genome-Scale CRISPR-Cas9 Knockout Screening in Human Cells. Science 343, 84–87 (2014).

76. Hart, T. et al. TKOv1 Library dataset. Cell (2015).

77. Hart, T., et al. TKOv3 Library Dataset. addgene (2017).

78. Release v2.0.0 - Supported libraries extended ⋅ francescojm/CRISPRcleanR,” GitHub. Available: /francescojm/CRISPRcleanR/releases/tag/2.0.0. [Accessed: Dec. 17, 2020]

79. “GRCh38.” Available: ftp://ftp.ncbi.nlm.nih.gov/genomes/archive/old_genbank/Eukaryotes/vertebrates_mammals/Homo_sapiens/GRCh38/seqs_for_alignment_pipelines/GCA_000001405.15_GRCh38_no_alt_analysis_set.fna.gz

80. Langmead, B., Trapnell, C., Pop, M. & Salzberg, S. L. Ultrafast and memory-efficient alignment of short DNA sequences to the human genome. Genome Biology 10, R25 (2009).

81. “GENCODE - Human Release 35.” Available: https://www.gencodegenes.org/human/release_35.html. [Accessed: Feb. 02, 2021]

82. Riester, L. W. and M. HGNChelper: Identify and Correct Invalid HGNC Human Gene Symbols and MGI Mouse Gene Symbols. (2019).

83. Oh, S. et al. HGNChelper: Identification and correction of invalid gene symbols for human and mouse. (2020) doi:10.12688/f1000research.28033.1.

84. Kim, E. & Hart, T. Improved analysis of CRISPR fitness screens and reduced off-target effects with the BAGEL2 gene essentiality classifier. Genome Medicine 13, 2 (2021).

85. “Hart-lab/bagel.” Hart Lab, Oct. 2020. Available: https://github.com/hart-lab/bagel. [Accessed: Dec. 17, 2020]

86. “Precaluclation of library alignment info (hart-lab/bagel),” GitHub. Available: https://github.com/hart-lab/bagel/blob/master/precalc_library_alignment_info.py. [Accessed: Feb. 02, 2021]

87. Kosty, J., Lu, F., Kupp, R., Mehta, S. & Lu, Q. R. Harnessing OLIG2 function in tumorigenicity and plasticity to target malignant gliomas. Cell Cycle 16, 1654–1660 (2017).

88. Hart, T., Brown, K. R., Sircoulomb, F., Rottapel, R. & Moffat, J. Measuring error rates in genomic perturbation screens: Gold standards for human functional genomics. Molecular Systems Biology 10, 733 (2014).

89. F. Iorio, “ADAM.” Jul. 2021. Available: https://github.com/francescojm/ADAM. [Accessed: Nov. 18, 2021]

90. Hart, T. & Moffat, J. BAGEL: A computational framework for identifying essential genes from pooled library screens. BMC bioinformatics 17, 164 (2016).

91. IntOGen - Cancer driver mutations in Glioblastoma.

92. McLaren, W. et al. The Ensembl Variant Effect Predictor. Genome Biology 17, 122 (2016).

93. Patel, A. P. et al. Single-cell RNA-seq highlights intratumoral heterogeneity in primary glioblastoma. Science (New York, N.Y.) 344, 1396–1401 (2014).

94. Müller, S. et al. A single-cell atlas of human glioblastoma reveals a single axis of phenotype in tumor-propagating cells. bioRxiv 377606 (2018) doi:10.1101/377606.

95. Gonzalez Castro, L. N. & Wesseling, P. The cIMPACT-NOW updates and their significance to current neuro-oncology practice. Neuro-Oncology Practice (2020) doi:10.1093/nop/npaa055.

96. Zhang, Y., Parmigiani, G. & Johnson, W. E. ComBat-seq: Batch effect adjustment for RNA-seq count data. NAR genomics and bioinformatics 2, lqaa078 (2020).

97. Robinson, M. D., McCarthy, D. J. & Smyth, G. K. edgeR: A Bioconductor package for differential expression analysis of digital gene expression data. Bioinformatics (Oxford, England) 26, 139–140 (2010).

98. Uno, M. et al. Correlation of MGMT promoter methylation status with gene and protein expression levels in glioblastoma. Clinics (Sao Paulo, Brazil) 66, 1747–1755 (2011).

99. Bell, R. J. A. et al. Cancer. The transcription factor GABP selectively binds and activates the mutant TERT promoter in cancer. Science (New York, N.Y.) 348, 1036–1039 (2015).

100. Bamshad, M. J. et al. Exome sequencing as a tool for Mendelian disease gene discovery. Nature Reviews Genetics 12, 745–755 (2011).

101. Yuan, X., Larsson, C. & Xu, D. Mechanisms underlying the activation of TERT transcription and telomerase activity in human cancer: Old actors and new players. Oncogene 38, 6172–6183 (2019).

102. Friedman, J. H., Hastie, T. & Tibshirani, R. Regularization Paths for Generalized Linear Models via Coordinate Descent. Journal of Statistical Software 33, 1–22 (2010).

103. Benkeser, D. & Van Der Laan, M. The Highly Adaptive Lasso Estimator. in 2016 IEEE International Conference on Data Science and Advanced Analytics (DSAA) 689–696 (2016). doi:10.1109/DSAA.2016.93.

104. Wright, M. N. & Ziegler, A. Ranger: A Fast Implementation of Random Forests for High Dimensional Data in C++ and R. Journal of Statistical Software 77, 1–17 (2017).

105. Chen, T. & Guestrin, C. XGBoost: A Scalable Tree Boosting System. in Proceedings of the 22nd ACM SIGKDD International Conference on Knowledge Discovery and Data Mining 785–794 (2016). doi:10.1145/2939672.2939785.

106. Coyle, J. Tmle3: The Extensible TMLE Framework. (2021) doi:10.5281/ZENODO.4603358.

107. laanvan der Laan, M. Finite Sample Inference for Targeted Learning. (2017) doi:10.48550/arXiv.1708.09502.

108. “HAL: The Highly Adaptive Lasso — fit_hal.” Available: https://tlverse.org/hal9001/reference/fit_hal.html. [Accessed: Nov. 09, 2022]

109. Shields, J. et al. VRK1 is a Novel Synthetic Lethal Target in VRK2-methylated Glioblastoma. 9.

110. Chapter 6 Super (Machine) Learning | Targeted Learning in R.

111. Wang, L. et al. The Phenotypes of Proliferating Glioblastoma Cells Reside on a Single Axis of Variation. Cancer Discovery 9, 1708–1719 (2019).

112. So, J. et al. VRK1 is required in VRK2-methylated cancers of the nervous system. 2021.12.28.474386 (2021) doi:10.1101/2021.12.28.474386.

113. Tamura, M. et al. PTEN interactions with focal adhesion kinase and suppression of the extracellular matrix-dependent phosphatidylinositol 3-kinase/Akt cell survival pathway. The Journal of Biological Chemistry 274, 20693–20703 (1999).

114. Thanapprapasr, D. et al. PTEN Expression as a Predictor of Response to Focal Adhesion Kinase Inhibition in Uterine Cancer. Molecular Cancer Therapeutics 14, 1466–1475 (2015).

115. Mohanty, A. et al. FAK-targeted and combination therapies for the treatment of cancer: An overview of phase I and II clinical trials. Expert Opinion on Investigational Drugs 29, 399–409 (2020).

116. Dawson, J. C., Serrels, A., Stupack, D. G., Schlaepfer, D. D. & Frame, M. C. Targeting FAK in anticancer combination therapies. Nature Reviews Cancer 21, 313–324 (2021).

117. Stafford, W. C. et al. Irreversible inhibition of cytosolic thioredoxin reductase 1 as a mechanistic basis for anticancer therapy. Science Translational Medicine 10, eaaf7444 (2018).

118. Auranofin reveals therapeutic anticancer potential by triggering distinct molecular cell death mechanisms and innate immunity in mutant P53 non-small cell lung cancer - ScienceDirect.

119. Wang, J. et al. Repurposing auranofin to treat TP53-mutated or PTEN-deleted refractory B-cell lymphoma. Blood Cancer Journal 9, 95 (2019).

120. Chen, L. et al. Novel inhibitors induce large conformational changes of GAB1 pleckstrin homology domain and kill breast cancer cells. PLoS computational biology 11, e1004021 (2015).

121. Gu, H. & Neel, B. G. The “Gab” in signal transduction. Trends in Cell Biology 13, 122–130 (2003).

122. Shalem, O. et al. GeckoV1 Library Dataset. Science (2014).

123. P. Paddison, “GSE70038.” Available: https://www.ncbi.nlm.nih.gov/geo/query/acc.cgi?acc=GSE70038. [Accessed: Dec. 21, 2020]

124. “Project Score Raw Readcounts.” Available: https://cog.sanger.ac.uk/cmp/download/raw_sgrnas_counts.zip

125. Dempster, J. M. et al. Extracting Biological Insights from the Project Achilles Genome-Scale CRISPR Screens in Cancer Cell Lines. bioRxiv 720243 (2019) doi:10.1101/720243.

126. Doench, J. G. et al. Optimized sgRNA design to maximize activity and minimize off-target effects of CRISPR-Cas9. Nature Biotechnology 34, 184–191 (2016).

127. DepMap. Achilles_common_essentials - DepMap 20q3.

128. Edinburgh Compute and Data Facility. The University of Edinburgh.

129. Kim, E. & Hart, T. Improved analysis of CRISPR fitness screens and reduced off-target effects with the BAGEL2 gene essentiality classifier. bioRxiv 2020.05.30.125526 (2020) doi:10.1101/2020.05.30.125526.

130. Xie, Y. Bookdown: Authoring Books and Technical Documents with R Markdown. (CRC Press, 2016).

131. Larsson, J. et al. Eulerr: Area-Proportional Euler and Venn Diagrams with Ellipses. (2021).

132. Vaughan, D., Dancho, M. & RStudio. Furrr: Apply Mapping Functions in Parallel using Futures. (2020).

133. Bengtsson, H. A unifying framework for parallel and distributed processing in r using futures. (2020).

134. “WebDriver,” *Selenium*. Available: https://www.selenium.dev/documentation/webdriver/. [Accessed: Jan. 28, 2022]

135. Köster, J. & Rahmann, S. Snakemake—a scalable bioinformatics workflow engine. Bioinformatics 28, 2520–2522 (2012).

136. Wickham, H. et al. Welcome to the Tidyverse. Journal of Open Source Software 4, 1686 (2019).

137. Coyle, J., Hejazi, N., Malenica, I., Sofrygin, O. & Phillips, R. Sl3: Modern Super Learning with Pipelines. (2021) doi:10.5281/ZENODO.1342293.

138. Huang, A., Garraway, L. A., Ashworth, A. & Weber, B. Synthetic lethality as an engine for cancer drug target discovery. Nature Reviews Drug Discovery 1–16 (2019) doi:10.1038/s41573-019-0046-z.

